# Identification of downstream effectors of retinoic acid specifying the zebrafish pancreas by integrative genomics

**DOI:** 10.1101/2020.11.09.341016

**Authors:** Ana R. López-Pérez, Piotr J. Balwierz, Boris Lenhard, Ferenc Muller, Fiona C. Wardle, Isabelle Manfroid, Marianne L. Voz, Bernard Peers

**Author notes:** Corresponding author: BP.

## Abstract

Retinoic acid (RA) is a key signal for the specification of the pancreas. Still, the gene regulatory cascade triggered by RA in the endoderm remains poorly characterized. In this study, we investigated this regulatory network in zebrafish by combining RNA-seq, RAR ChIP-seq and ATAC-seq assays. By analysing the effect of RA and of the RA receptor (RAR) antagonist BMS439 on the transcriptome and on the chromatin accessibility of endodermal cells, we identified a large set of genes and regulatory regions regulated by RA signalling. RAR ChIP-seq further defined the direct RAR target genes including the known *hox* genes as well as several pancreatic regulators like *mnx1, insm1b, hnf1ba* and *gata6*. Comparison of our zebrafish data with available murine RAR ChIP-seq data highlighted conserved direct target genes and revealed that some RAR sites are under strong evolutionary constraints. Among them, a novel highly conserved RAR-induced enhancer was identified downstream of the *HoxB* locus and driving expression in the nervous system and in the gut in a RA-dependant manner. Finally, ATAC-seq data unveiled the role of the RAR-direct targets Hnf1ba and Gata6 in opening chromatin at many regulatory loci upon RA treatment.

**Summary statement:** Combination of RNA-seq, ChIP-seq and ATAC-seq assays identifies genes directly and indirectly regulated by RA signalling in zebrafish endoderm. Comparison with murine data highlights RAR binding sites conserved among vertebrates.

## Introduction

Retinoic acid (RA) is essential for the development of vertebrate embryos. This small lipophilic molecule controls the formation of many organs and tissues and is crucial for the anteroposterior (AP) patterning of the hindbrain and endoderm (Rhinn and Dollé 2012; Niederreither and Dollé 2008; Ghyselinck and Duester 2019). During gastrulation and early somitogenesis, RA forms a two-tailed gradient with highest levels in the mid-trunk region and progressive lower levels at the anterior and posterior regions of the embryo (Shimozono et al. 2013). This gradient regionalizes the endoderm along the AP axis and specifies distinct organs at the appropriate location (Larsen and Grapin-Botton 2017; Zorn and Wells 2009). RA is notably required for the specification of the pancreatic field in several animal models like zebrafish, xenopus, chick and mice (David Stafford and Prince 2002; D. Stafford et al. 2004; Bayha et al. 2009; Chen et al. 2004; Molotkov, Molotkova, and Duester 2005; Martín et al. 2005) and protocols for generating pancreatic cells *in vitro* from embryonic stem cells include a RA incubation step of endodermal cells (Pagliuca et al. 2014). Experiments in zebrafish have shown that, when RA signalling is blocked pharmacologically by incubating embryos with the pan-RAR antagonist BMS493, no pancreas develops; conversely, when embryos are incubated with exogenous RA, ectopic pancreatic cells are generated in the anterior endoderm. Such treatments performed at different stages revealed that RA instructs the endoderm to form pancreas during late gastrulation (from about 8 to 12 hours post-fertilization, hpf)(David Stafford and Prince 2002). Cell transplantation studies have also shown that RA, synthesized in the anterior paraxial mesoderm, acts directly on the mid-trunk endoderm to promote pancreas development (David Stafford et al. 2006). Although several regulatory genes have been identified as important downstream targets of RA, such as *Hox* or *Hnf1b* genes (Song et al. 2007; D. Stafford et al. 2004; Hernandez et al. 2004; Nolte, De Kumar, and Krumlauf 2019; Gere-Becker et al. 2018), the regulatory network triggered by RA in endoderm during gastrulation is still unclear.

RA levels are tightly controlled by the balance of its synthesis, controlled mainly by Aldh1a2 (Raldh2), and its degradation catalysed by Cyp26a enzymes (Niederreither and Dollé 2008). This synthesis/degradation balance is notably regulated by auto-regulatory loops where RA levels control expression of some of its metabolizing enzymes (e.g. *Cyp26a* and *Dhrs3*)(Kinkel et al. 2009; Feng et al. 2010a). RA controls gene expression through binding to Retinoic Acid Receptors (RAR) which form heterodimers with Retinoid X Receptors (RXR). These RAR/RXR heterodimers bind to DNA regulatory sequences, called “Retinoic Acid Response Elements” (RAREs), located in promoters or enhancers. Genome-wide identification of RAR binding sites has been achieved by ChIP-seq experiments mostly using murine or human cell lines (Delacroix et al. 2010; Mahony et al. 2011; Chatagnon et al. 2015; Moutier et al. 2012). Such studies confirmed that the RAR/RXR heterodimers bind to direct repeats of the RGKTCA motif (R=A/G, K=G/T) usually separated by 5 bases (DR5), but also to repeats of this motif with other spacing and orientations. These data uncovered several thousand RAR/RXR binding sites in the murine and human genomes, some located near RA-induced genes like the *Hox, Cyp26a*, or *Rar* genes. However, it is not yet clear if all the identified RAR binding sites have a regulatory function.

In order to decipher the gene regulatory network triggered by RA during gastrulation in zebrafish endodermal cells, we combined RNA-seq, ChIP-seq and ATAC-seq approaches. We identified genes regulated by RA signalling by analysing the transcriptome of endodermal cells from RA- or BMS493-treated embryos. Direct targets of RAR were identified by ChIP-seq assays. Then, ATAC-seq experiments (Buenrostro et al. 2013) allowed us to identify chromatin regions whose accessibility is modified upon RA treatment. By integrating all these data, we identify important transcription factors acting downstream RA signalling in endodermal cells and involved in pancreas development. Furthermore, comparison of the RAR sites detected in the zebrafish genome and in the murine genome unveiled RAR target genes which have been maintained during evolution and whose RAR site sequences are well conserved. By this approach, RAR sites with essential function can be identified in vertebrate genomes.

## Results

### Retinoic acid affects the transcriptome of zebrafish endodermal cells

To identify all genes regulated during gastrulation by RA in zebrafish endodermal cells, transgenic Tg(*sox17*:GFP) zebrafish gastrulae were treated either with 1µM RA, with 1µM BMS493 (pan-RAR-antagonist) or with DMSO (control). Endodermal GFP+ cells were next selected by FACS from embryos at 3-somites (3-S) and 8-somites (8-S) stages (11-hpf and 13.5-hpf, respectively). Non-endodermal (GFP-) cells were also selected from the DMSO-treated control embryos in order to compare with GFP+ endodermal cells and identify genes displaying endodermal enriched expression. RNA-seq was performed on all these FACS-isolated cells prepared in triplicate (24 samples in total) and transcriptomes were analysed using the bioinformatic pipeline as described in Methods. Principal component analysis of all RNA-seq data (Fig.1A) shows i) a tight clustering of all triplicate samples confirming a high reproducibility in the experiment, ii) a strong difference between the transcriptome of endodermal and non-endodermal (NE) cells (discriminated along the first axis of the PCA plot), iii) relatively similar transcriptomes of cells isolated at 3-S and 8-S stages, and iv) a clustering of BMS493 samples near DMSO samples indicating a much weaker effect of BMS493 treatments compared to the RA treatments. These conclusions were further confirmed by the differential gene expression analyses between the different conditions. Indeed, more differentially expressed genes were identified between endodermal and non-endodermal cells (1370 and 1410 differentially expressed genes at 3-S and 8-S stages, respectively with a FDR<0.01) than between endodermal cells treated with RA versus DMSO (756 and 514 RA-regulated genes at 3-S and 8-S stages, respectively with FDR<0.01) or with BMS493 versus DMSO (32 and 71 BMS493-regulated genes at 3-S and 8-S stages, respectively). We found a large overlap among the sets of genes having an endodermal-enriched expression at 3-S and at 8-S stages (Fig. S1A; list of genes given in Table S1) and these sets include all known endodermal markers including *sox17, gata5/6* and *foxa1/2/3*, validating the accurate sorting of endodermal cells. RA-regulated genes consist of a large set of up- and down-regulated genes (Table S2), many of them being regulated at both 3-S and 8-S stages (Fig. S1B). Some of these RA-regulated genes correspond to known RAR-direct targets such as *cyp26b1/a1, dhrs3a* and several *hox* genes, confirming the efficiency of RA treatments. Interestingly, BMS493 treatment led mostly to down-regulation of gene expression. Indeed at 3-S stage, the 32 BMS493-regulated genes were all repressed, and, at 8-S stage, 68 genes were repressed while only 3 genes were up-regulated by BMS493 treatment (Tables S3). A large overlap was also observed between the genes down-regulated by BMS493 at 3-S and at 8-S stage (Fig. S1C). As expected, a large proportion of genes down-regulated by BMS493 were up-regulated by RA treatment, this observation being evident mostly at 3-S stage where 72% of genes repressed by the RAR antagonist were induced by RA, while this proportion decreased to 40% at 8-S stage (Fig. 1B and C). Tables 1 and 2 show the genes significantly up-regulated by RA and down-regulated by BMS493 as well as those enriched in the endoderm at 3- and 8-somites stage, respectively (shown in green). *gata6, insm1a* and *ascl1b* are the only known pancreatic regulatory genes which were regulated by both RA and BMS493 (Table 1). Other pancreatic transcription factors, like *mnx1, insm1b, hnf1ba, nr5a2 o*r *neuroD1*, were induced by the RA-treatment but not significantly repressed by BMS493. Inversely, other pancreatic regulators were inhibited by BMS493 but not significantly induced by RA like *pdx1, rfx6* and *myt1b* (Tables S3 and S6). We can assume that the induction of pancreatic fate by RA (and the absence of pancreas upon BMS439 treatment) is mediated, at least in part, by the direct or indirect regulation of these pancreatic regulatory factors. In conclusion, these RNA-seq data highlight all the genes with an enriched expression in zebrafish endodermal cells and which are regulated by RA signalling. This gene set notably includes regulators involved in the AP patterning of the endoderm, such as *hox* genes, and known factors involved in the specification of pancreatic progenitors.

**Table 1:**
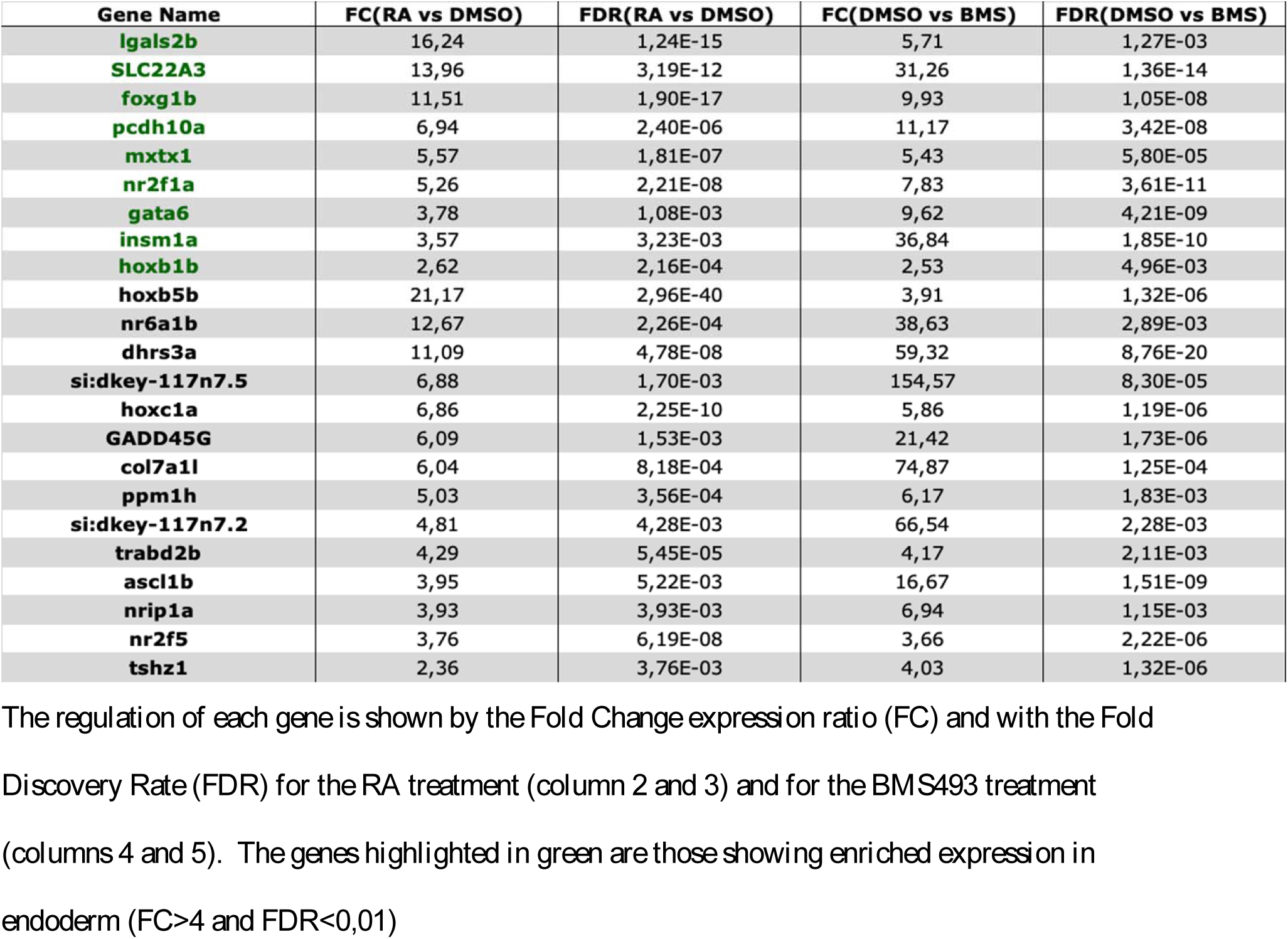
List of genes up-regulated by RA and down-regulated by BMS493 at 3-somites stage (with FDR<0,01).

**Table 2:** List of genes up-regulated by RA and down-regulated by BMS493 at 8-somites stage (with FDR<0,01).

| Gene Name | FC(RA vs DMSO) | FDR(RA vs DMSO) | FC(DMSO vs BMS) | FDR(DMSO vs BMS) |
| --- | --- | --- | --- | --- |
| **hoxb5b \*** | 12,07 | 1,55E-32 | 6,26 | 7,24E-16 |
| **slc13a2** | 8,58 | 2,14E-07 | 11,18 | 7,32E-07 |
| **dhrs3a \*** | 8,13 | 2,40E-18 | 100,51 | 1,57E-56 |
| **hoxb6b** | 7,46 | 4,15E-10 | 3,40 | 5,40E-03 |
| **si:ch211-157b11.8** | 7,65 | 1,62E-06 | 5,32 | 3,25E-03 |
| **SLC22A3 \*** | 5,43 | 1,09E-08 | 30,78 | 1,85E-27 |
| **ezra** | 5,25 | 4,97E-04 | 7,89 | 3,59E-05 |
| **tmem30c** | 5,15 | 9,33E-05 | 8,01 | 1,85E-05 |
| **cldn15a** | 4,66 | 7,70E-04 | 10,12 | 1,86E-07 |
| **pcdh10a \*** | 4,41 | 2,42E-04 | 9,77 | 8,08E-09 |
| **epb41l4a** | 3,57 | 6,96E-06 | 7,06 | 6,52E-11 |
| **nr2f1a \*** | 3,53 | 4,28E-05 | 5,98 | 3,93E-09 |
| **hmga2** | 2,54 | 5,11E-04 | 3,18 | 6,39E-05 |
| **hoxa3a** | 18,66 | 9,70E-40 | 3,41 | 1,59E-05 |
| **skap1** | 12,49 | 8,28E-17 | 5,17 | 1,66E-04 |
| **masp1** | 9,61 | 2,19E-10 | 17,11 | 2,99E-07 |
| **foxg1b \*** | 8,27 | 1,11E-06 | 25,02 | 2,99E-07 |
| **si:dkey-172f14.2** | 8,00 | 7,40E-08 | 14,27 | 1,77E-04 |
| **zgc:194209** | 7,70 | 1,29E-07 | 6,84 | 6,66E-04 |
| **cldn2** | 7,40 | 2,18E-06 | 11,58 | 1,85E-05 |
| **si:dkey-25e11.10** | 5,68 | 8,30E-06 | 6,95 | 1,05E-03 |
| **ascl1a** | 5,61 | 7,72E-08 | 11,93 | 2,35E-07 |
| **cldnc** | 5,19 | 1,96E-06 | 10,72 | 8,81E-12 |
| **hoxc1a \*** | 5,00 | 1,10E-05 | 5,90 | 1,09E-05 |
| **iqgap2** | 4,75 | 1,55E-04 | 4,80 | 3,76E-03 |
| **zgc:113054** | 2,90 | 1,66E-04 | 3,36 | 8,39E-05 |
| **tshz1 \*** | 2,90 | 1,51E-03 | 2,93 | 7,29E-03 |

**Figure 1.**
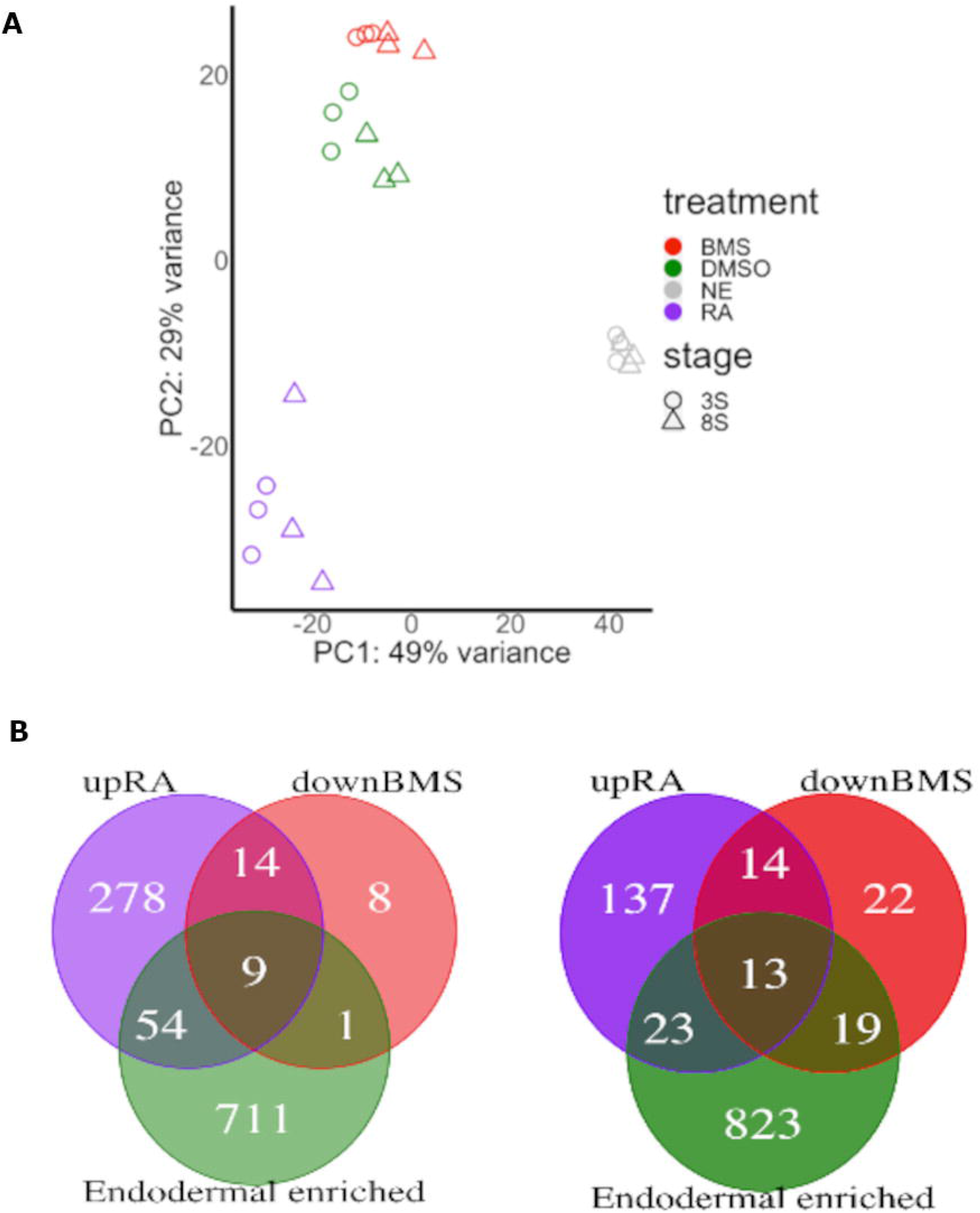
Effect of RA and BMS493 on the transcriptome of endodermal cells. (A) Principle component analysis (PCA) of the 24 RNA-seq data obtained on cells isolated at 3-somites stage (circle) and 8-somites stage (triangle). The colors indicate the data for non-endodermal cells (NE)(grey), and endodermal cells treated with RA (purple), BMS493 (red) and DMSO as control (green). The plot shows high reproducibility between triplicates. The strongest transcriptomic differences occur between endodermal and non-endodermal cell (along PC1), then between endodermal cells treated with RA and DMSO (along PC2), while BMS493 treatment has minimal influences. Consistent with the antagonist action of BMS493 and with the agonist action of RA, these samples are located far from each other, the DMSO-samples being located between them. (B-C) Venn diagram displaying the number of genes up-regulated by RA (purple), down-regulated by BMS493 (red) and endodermal enriched (green) at 3-somites (B) and 8-somites stage (C).

### Identification of RAR binding sites in the zebrafish genome

To further identify the genes directly regulated by RAR and determine if some pancreatic regulatory genes are direct targets of RA signalling, we performed ChIP-seq experiments at the end of gastrulation. In absence of validated commercialized ChIP-grade antibody recognizing zebrafish RAR, we chose to express a tagged RARaa in zebrafish gastrulae by injecting zebrafish fertilized eggs with the mRNA coding for the zebrafish RARaa protein fused to a Myc-tag at its C-terminal end. RARaa was chosen as the RNA-seq data indicated that it is the most highly expressed RA receptor in zebrafish endodermal cells. Injection of this mRNA did not disturb the development of the embryos. Chromatin was prepared at 11.5 hpf (3 somites stage) from about 2000 injected zebrafish embryos and immunoprecipitation was performed with a ChIP-grade Myc antibody. Comparison of reads obtained with the Myc-RAR ChIP and the input negative control led to the identification of 4858 RAR peaks. In order to identify *bona fide* RAR binding sites showing strong affinity, we selected all peaks with a height score above 50 (Table S4). By choosing such criteria, 2848 robust RAR binding sites were identified in the zebrafish genome. As shown in Figure 2A, a majority of these sites are located near or within genes: 8% were found in promoters (i.e. 1kb upstream of the gene TSS, Transcription Start Site), 30% in upstream sequences (from 1 to 10 kb), 22% in introns, 4% in exons, and only 33% in intergenic regions. Sequence analysis of all RAR peaks revealed that the highest represented motif corresponds to the <u>D</u>irect <u>R</u>epeat of the RGKTCA motif separated by <u>5</u> bases (reported as DR5) and being the RAR/RXR consensus binding sequence (present in 39% of identified RAR peaks)(Fig. 2B)(Rhinn and Dollé 2012). The next most abundant motifs are also repetitions of the RGKTCA motif with different spacings (TR4 being a <u>D</u>irect <u>R</u>epeat separated by 1 and 2 bases: DR1/DR2) and orientations (Rxra being an <u>I</u>nverted <u>R</u>epeat with no base separation: IR0). Furthermore, many RAR peaks were found near genes known to be RAR-direct target genes, such as *cyp26a1, dhrs3* or the *hoxb1a-hoxb4a* genomic region (Fig. 2C and data not shown). All these observations confirm the accuracy of the ChIP-seq data. Interestingly, many identified zebrafish RAR sites are located in evolutionary conserved genomic sequences as shown by the fish PhastCons track (Fig. 2C and see below).

**Figure 2.**
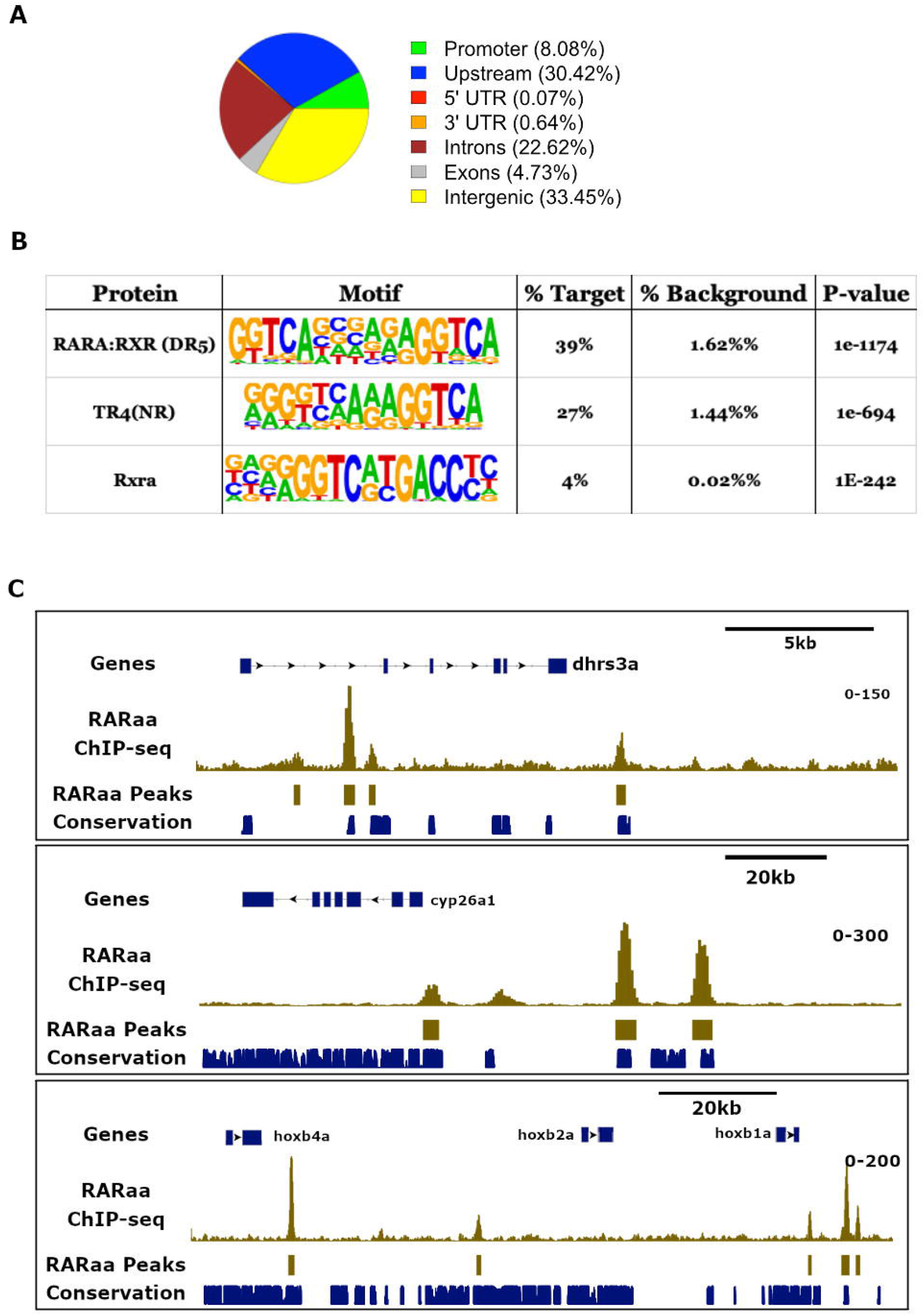
Identification of RAR binding sites in the zebrafish endoderm. (A) Distribution of ChIP-seq peaks to the different regions of the zebrafish genome. (B) Top 3 motifs overrepresented in all ChIP-seq peak sequences with the percentage of sites containing the motifs and the p-value of enrichment. The three motifs consist to a repetition of the A/GGGTCA sequence; the first corresponds to the classical DR5 recognized by the RARA:RXR complex, the second (TR4) is a superposition of DR1 and DR2, and the third is a IR0. (C) Visualization of RARaa binding sites around the *dhrs3a* gene (upper panel), *cyp26a1* (middle panel) and *hoxb1a-hoxb4a* genomic region (below panel). Tracks in gold correspond to RARaa ChIP-seq reads and identified RARaa peaks. The track in blue shows the location of conserved genomic sequences (from the UCSC Genome Browser obtained from comparison of 5 fish species).

The RAR ChIP-seq peaks located within 250 kb from the TSS of a gene were assigned to this gene and when several genes were lying in the vicinity of a RAR site, the closest gene was considered as the putative RAR-regulated gene. Using this strategy, amongst the 2848 RAR sites, 2144 were linked to a gene. Correlation analysis between RA gene expression regulated by RA and the number of RAR sites near the gene showed that RAR acts mainly as a transcriptional activator (Fig. 3A), supporting the classical model where RAR/RXR heterodimers mainly recruit co-activators upon RA ligand binding (Rhinn and Dollé 2012). However, this correlation is not very high and genes down-regulated by RA can harbour nearby RAR sites. Indeed, from the gene set having RAR sites, 61 were down-regulated and 94 genes were up-regulated by RA at 3-S stage (Fig. 3B, Tables S5). Among the genes up-regulated by RA and harbouring a RAR site, we identified the pancreatic regulatory genes *hnf1ba/b, gata6, insm1b, jag2b* and *mnx1* indicating that these genes are direct targets of RA. As for the genes down-regulated by BMS493, we found that a large number of them (i.e. 18 genes out of 32) contain RAR sites and amongst them 13 are also upregulated by RA (Fig. 3B, see legends for gene names). In conclusion, the ChIP-seq data allowed us to define the zebrafish RAR cistrome at the end of gastrulation and identify putative RAR direct target genes.

**Figure 3.**
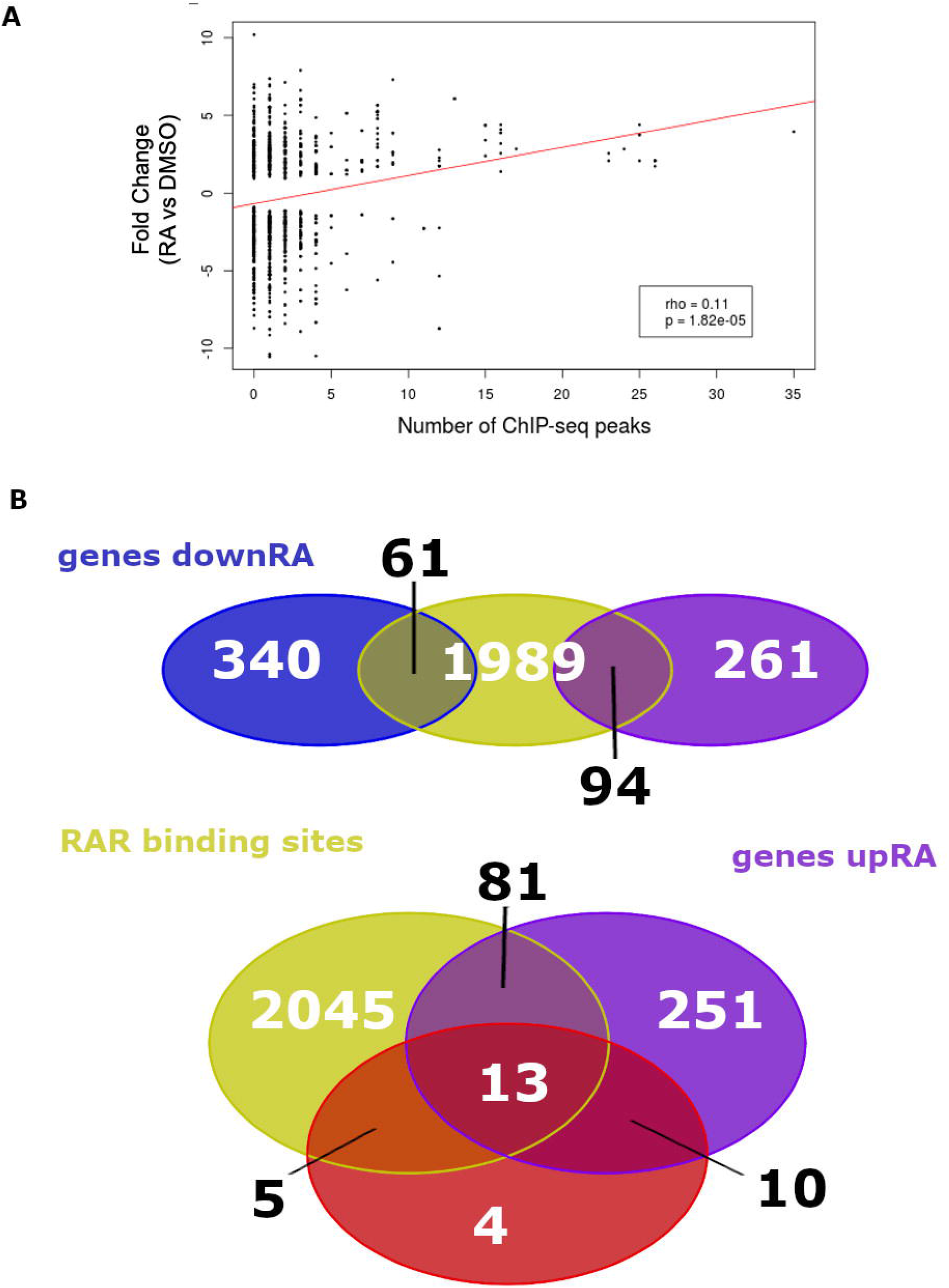
Integrated analysis of the ChIP-seq and RNA-seq data. (A) Correlation of RA gene regulation (log2 fold change of expression RA versus DMSO) according to the number of neighbouring RARaa ChIP-seq peaks. Only RA-regulated genes were included in the plot. (B) Venn diagram showing the overlap of genes harbouring nearby RARa binding sites (yellow) and those up-regulated by RA (purple), or down-regulated by RA (blue). The below panel also shows the overlap with the genes down-regulated by BMS493 (red). The 13 genes showing up-regulation by RA, down-regulation by BMS493 and harbouring a RAR site are *tshz1, nr6a1b, foxg1b, nr2f5, gata6, dhrs3a, hoxb1b, slc22a3, ppm1h, nrip1a, col7a1l, hoxc1a* and *hoxb5b*.

### Conservation of RAR binding sites from fish to mammals

Functionally important regulatory regions are expected to be conserved during evolution. To determine which RAR binding sites are conserved in vertebrates, we compared our zebrafish ChIP-seq data with those of Chatagon and colleagues (Chatagnon et al. 2015) who identified RAR binding sites in the murine genome using the F9 embryonal carcinoma cells whose differentiation into primitive endodermal cell is induced by RA treatment. This comparison revealed that, among the 2144 zebrafish genes harbouring a RAR site, 722 have also a RAR site near the murine orthologous genes. This list of conserved RAR-bound genes comprises notably *cyp26a1/b1, dhrs3a*, many *hox* genes, *raraa/b* as well as pancreatic genes *gata6, hnf1ba, insm1* and *mnx1*. We next determined which of these RAR binding sites are located in conserved regulatory sequences. To that end, we retrieved the list of conserved non-coding elements (CNEs) identified in zebrafish by comparing multiple fish and tetrapod genomes (Hiller et al. 2013). Amongst the 722 conserved RAR-bound genes, 116 RAR binding sites were located in CNEs, supporting a regulatory function (Table S6). Amongst them, 24 CNEs were even conserved from fish to mouse and are called here HCNE for highly conserved non-coding elements. They are found for example near the *meis1/2, srsf6, qki, nrip1* and *ncoa3* genes (Table S6). As already reported, several RAR sites controlling the expression *of hox* genes are located in these HCNEs (Nolte, De Kumar, and Krumlauf 2019). Interestingly, we identified here a novel RAR site in a HCNE located 25 kb downstream of the zebrafish *hoxb* cluster in the fourth intron of the *skap1* gene. This intron contains 4 strong RAR sites in the zebrafish and murine sequences (green boxes in Fig. 4A and B) and the sequence of the second RAR site has been maintained throughout vertebrate evolution and shows a motif similar to a DR5 RARE (Fig 4C). The transcriptional regulatory function of this HCNE was tested by transgenesis by inserting one copy of this element upstream of a minimal *cfos* promoter driving GFP. As shown in Fig 5A, this reporter transgene is expressed in the gut and in the neural tube. Highest GFP levels were detected in the posterior hindbrain, in a pattern highly reminiscent of *hoxb1b* gene expression. Furthermore, when transgenic embryos were treated with exogenous RA, GFP expression was drastically increased and detected in the whole morphologically affected embryos (Fig. 5B). Conversely, treatments with the RA antagonist BMS493 turned off the expression of this DR5-RAR-skap1:GFP transgene (Fig. 5C). This confirms the RARE function of this element and indicates a role in *hoxbb* cluster regulation.

**Figure 4.**
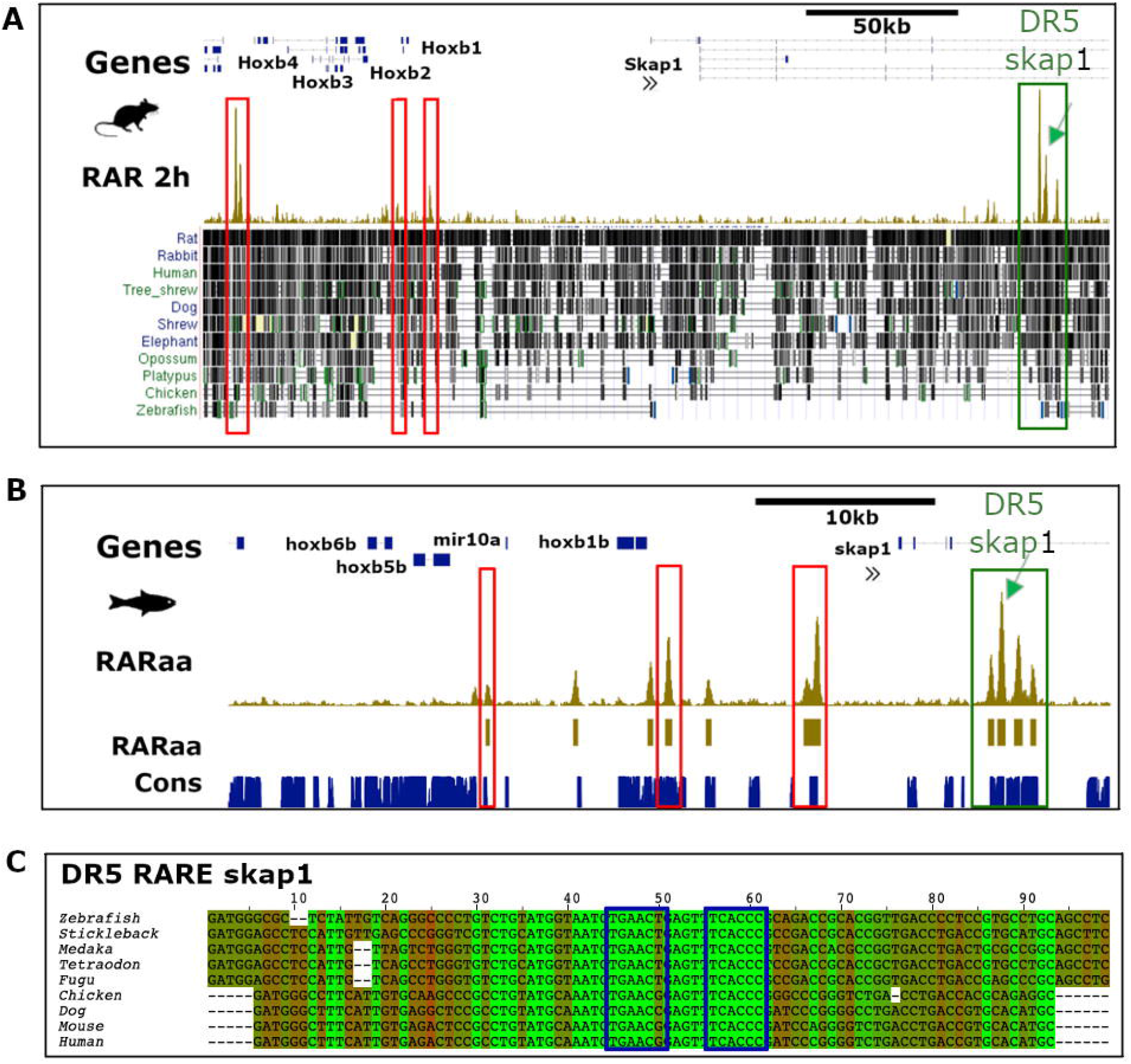
Examples of RARa binding sites conserved among vertebrates. (A-B) Genome browser views around HoxB-Skap1 locus in mouse (A) and zebrafish (B) showing the RAR binding sites detected by ChIP-seq (in gold) in both species. All RAR sites are located in CNEs but only the RAR sites located in skap1 gene 4th intron (green box) display sequence conservation from zebrafish to human. Other RAR sites located at similar places in the murine and zebrafish loci (red boxes) are probable orthologous RAR sites but their sequences cannot be aligned between zebrafish and mice. Conserved regions from the UCSC genome browser is shown below the murine RAR ChIP-seq peaks in panel A. (C) Alignment of 9 vertebrate sequences corresponding to the second RAR sites located in skap1 gene 4^th^ intron, showing the conservation of a DR5-like motif (blue boxes; inverse orientation) recognized by the RAR-RXR complex. Sequences highlighted in green are identical in all 9 species.

**Figure 5.**
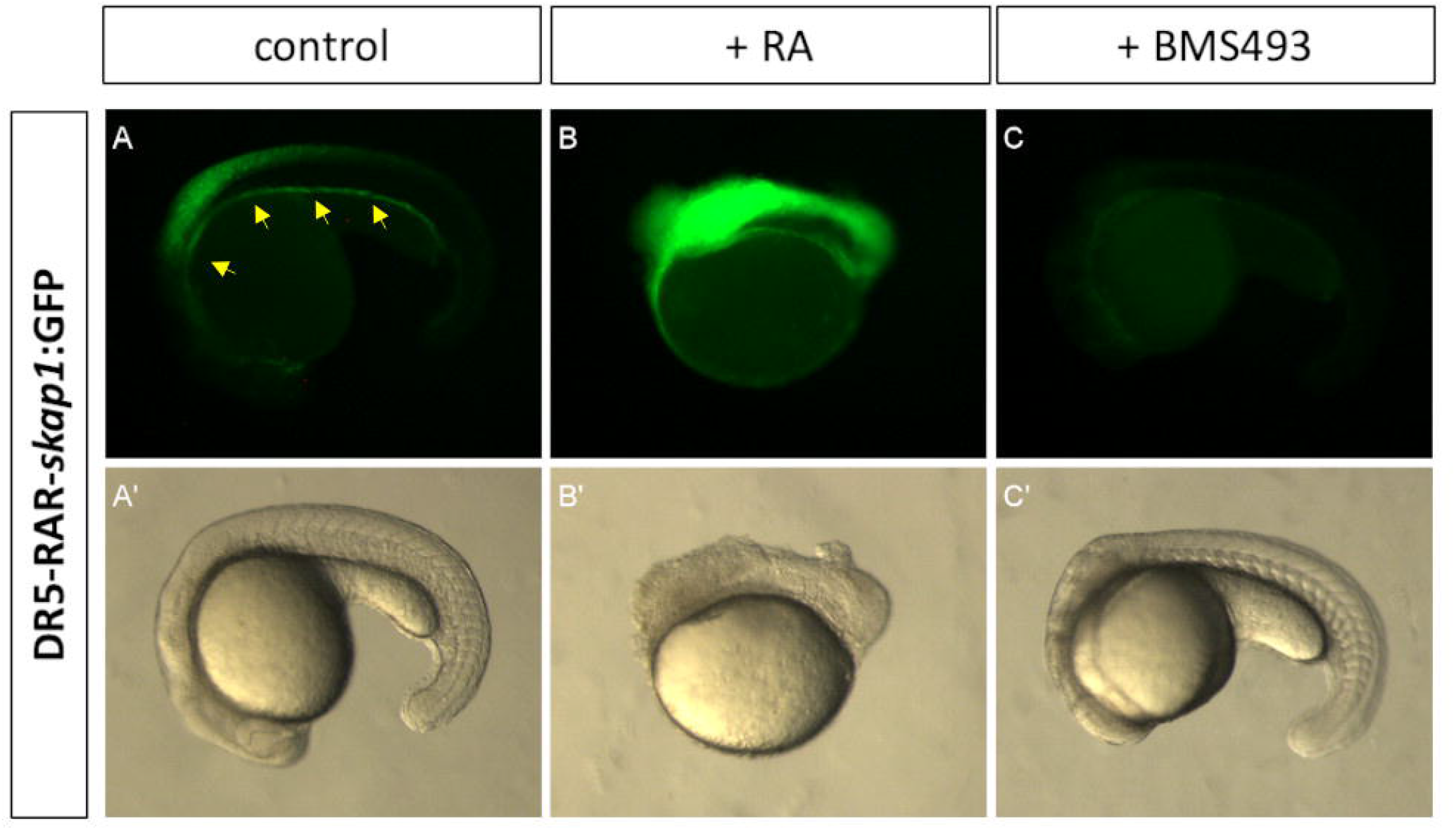
The conserved RAR site from skap1 gene 4th intron is a functional RARE. Pictures of the DR5-RAR-skap1:GFP transgenic embryos treated with DMSO (control, panels A), with RA (panels B) or with BMS493 (panels C). Upper panels display GFP fluorescent expression and lower panels (A’, B’ and C’) display embryo morphology. GFP Expression in the posterior endoderm gut is indicated by yellow arrows.

We also verified the regulatory role of some other RAR sites identified near zebrafish pancreatic regulatory genes. For example, we tested the activity of a RAR site located near a CNE downstream of the zebrafish *mnx1* gene. A RAR binding site is also found at a similar location downstream of the murine *Mnx1* gene (Fig.S2A). This RAR element was able to target GFP expression to the posterior endoderm at low level and in the dorsal pancreatic bud at higher levels (Fig S2 B-D). When the RAR-*mnx1*:GFP transgenic embryos were treated with RA, GFP expression was slightly increased in whole endoderm; conversely, treatment with BMS493 abolished GFP expression (Fig S2 D-F).

In conclusion, these analyses show that a large fraction of RAR sites has been conserved during evolution and transgenic assays confirm that some elements are sufficient to drive expression in endoderm and confer a RA-response.

### RA affects chromatin accessibility in endodermal cells

RAR-RXR complexes control gene expression through the recruitment of corepressors (e.g. NCoR/Smart) and coactivators (NCoA) which control chromatin compaction via HDACs and HATs, respectively (Ghyselinck and Duester 2019; Niederreither and Dollé 2008). Thus, one possible strategy to identify functional RAREs is to perform a genome-wide analysis of chromatin accessibility modifications following RA or RA-antagonist treatments by ATAC-seq assays (Buenrostro et al. 2013) allowing the identification of open chromatin and nucleosome-free regions induced or repressed by RA signaling. As most nucleosome-free regions correspond to regulatory sequences, ATAC-seq can also highlight the enhancers or promoters whose accessibility is modified by RA signalling indirectly, i.e. through the binding of transcription factors whose expression is induced by RA. Thus, sequence analysis of all RA-induced ATAC-seq peaks can give clues on the identity of transcription factors acting in the subsequent steps of the RA-induced regulatory cascade.

As for the RNA-seq assays, zebrafish embryos were treated with RA, BMS493 and DMSO during gastrulation and about 10000 endodermal cells were selected by FACS at 3-somites stage (11hpf). Non-endodermal cells from control DMSO-treated embryos were also analysed in parallel. Cell preparations and ATAC-seq experiments were done in triplicate for each condition and analysed as described in Methods. We first verified the accuracy of the data by several quality control analyses. First, for all samples, the analysis of the ATAC-seq fragment size distribution revealed the expected pattern with abundant short (<150 bp) fragments corresponding to nucleosome-free regions and larger fragments of about 200 and 400 bp corresponding to mono- and bi-nucleosome regions, respectively (Fig. S3A). Secondly, as reported previously (Quillien et al. 2017; Buenrostro et al. 2013), genome mapping of the nucleosome-free fragments showed a clear enrichment in promoter regions immediately upstream of transcriptional start sites (TSSs), while mono-nucleosomes were depleted from TSSs and rather mapped just downstream of the TSSs in a periodic manner (Fig. S3B). Thirdly, we verified that the ATAC-seq fragments correspond to many zebrafish regulatory regions by comparing them with regions harbouring the histone modifications H3K4me3 and H3K27ac marks identified in zebrafish embryos at 10 hpf (Paik et al. 2013). Heatmap plots of ATAC-seq reads from all samples showed an obvious enrichment at loci harbouring these two histone modifications (Fig. 6A and Fig. S4). As regulatory regions often display sequence conservation, we also compared our ATAC-seq reads to the collection of zebrafish evolutionary-conserved non-coding elements (zCNEs)(Hiller et al. 2013). Heat-maps of ATAC-seq reads from each sample also showed a strong correlation with zCNEs (Fig. 6A and Fig. S4). These observations confirm that regions identified by ATAC-seq exhibited hallmarks of active regulatory elements. The reproducibility of ATAC-seq data was also analysed by PCA (Fig. 6B). This PCA showed that i) triplicate samples are tightly clustered, and ii) endodermal and non-endodermal (NE) cell clusters are separated along the PC2 axis, while the RA-treated cluster is separated from the DMSO- and BMS-cell cluster along the PC3 axis. Thus, as observed for the RNA-seq data (Fig.1A), stronger differences are observed between GFP+ and GFP-cells compared to the differences between RA-treated and DMSO-treated endodermal cells. The samples corresponding to the BMS493-treated cells and DMSO-treated cells were not clearly segregated and did not reveal significant effects of BMS493 on the chromatin accessibility.

**Figure 6.**
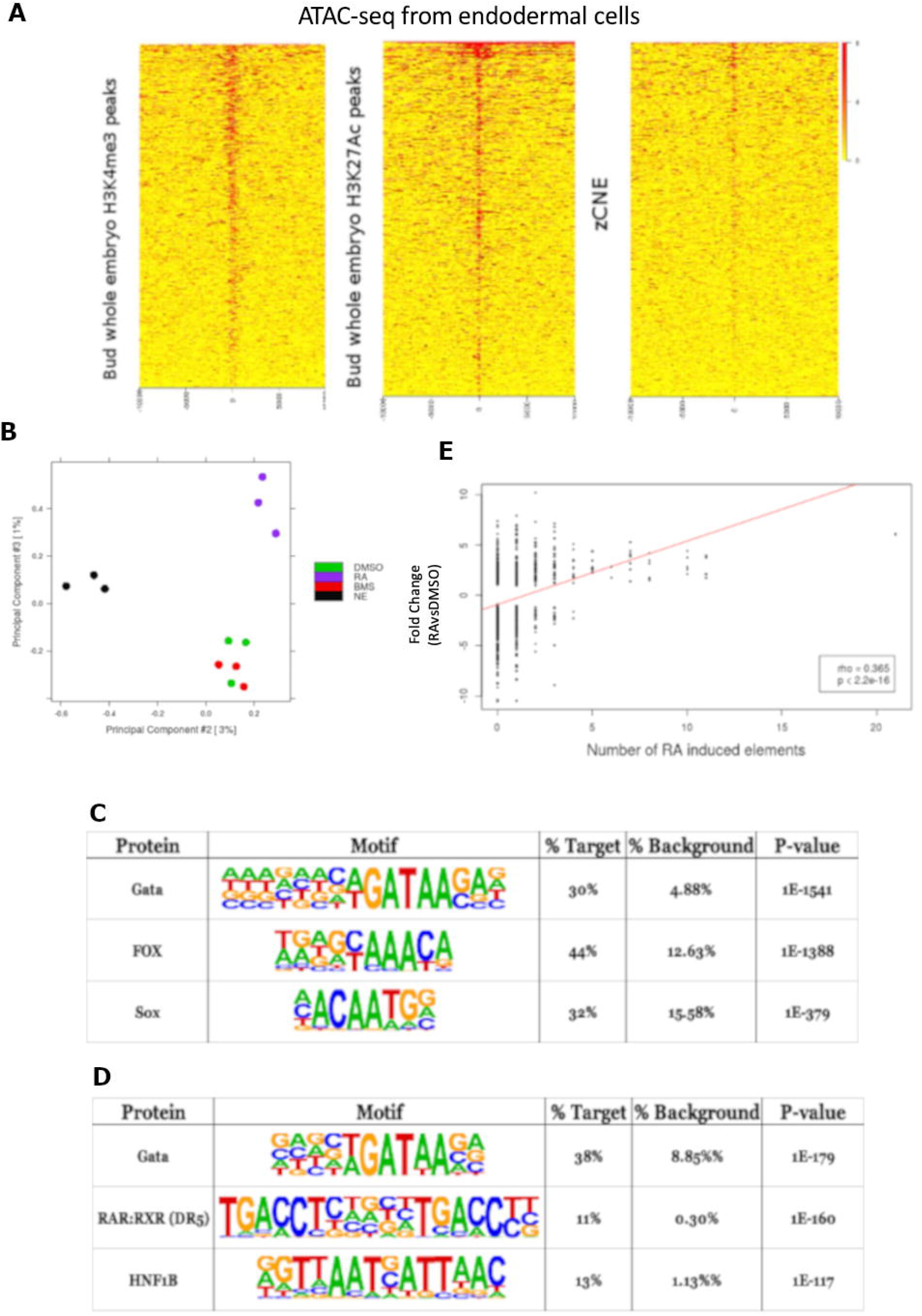
Identification of nucleosome-free regions in zebrafish endodermal cells and following RA treatments by ATAC-seq assays. (A) Heat maps showing enrichment of ATAC-seq reads at the middle of chromatin regions harbouring H3K4me3 and H3K27Ac epigenetic marks and at genomic areas corresponding to zCNE. The maps display intervals flanking 10 kb up and downstream of the features. The heat map plots shown on this figure corresponds to the ATAC-seq data obtained with control endodermal cells (DMSO-treated). Similar results were obtained for the other samples (see Suppl. Figure 4). (B) PCA plots obtained for the ATAC-seq libraries. ATAC-seq data from endodermal and non-endodermal cells are separated along the PC2 axis, while those from RA-treated versus control and BMS493 are separated along the PC3 axis. The plot shows clustering of triplicates and no obvious separation of the DMSO- and BMS493-treated cells. (C-D) Top 3 enriched motifs found in nucleosome-free regions detected specifically in endoderm (C) and detected following RA-treatments (D). (E) Plot showing the correlation of RA-induced gene expression (log2 fold change) and the number of RA-induced nucleosome-free elements located nearby the genes. Only genes showing significant RA gene induction were included.

From all 12 ATAC-seq samples, a total of 156,604 nucleosome-free regions were identified within the zebrafish genome. Differential peak intensity analyses revealed that 9722 and 12974 regions are more accessible in endodermal and non-endodermal cells, respectively (with FDR<0.05, Tables S7). Interestingly, sequence analysis of all endodermal-specific ATAC-seq regions highlight a significant enrichment of the binding site motifs for the Gata, Fox and Sox protein families (Fig. 6C), in accordance with the well-known function of Gata4/5/6, Foxa1/2/3 and Sox32/17 in endodermal cell differentiation (Zorn and Wells 2009). Also, these endoderm-enriched ATAC-seq peaks are often identified near endodermal and pancreatic regulatory genes, such as in the *foxa2, nkx6*.*1, hnf4g, sox17* or *mnx1* loci (Fig. S5 and data not shown), suggesting the presence of endoderm-specific enhancers at these locations.

As expected, the RA treatments had less influence on ATAC-seq peaks compared to the cell type identity (i.e. endoderm versus non-endoderm). Still, 1240 genomic regions were found to be more accessible and 749 regions were less accessible in the RA-treated endoderm compared to the DMSO-treated controls (with FDR<0.05). If the RA-treated samples were directly compared with the BMS493-treated samples, more peaks were detected as treatment-dependent: 3277 regions were more accessible in RA-treated while 1762 were more accessible in the antagonist-treated cells (Tables S8). Annotation of the RA-induced ATAC-seq peaks to the closest gene revealed that they were often located near RA-upregulated genes identified above by RNA-seq. Moreover, we found a significant correlation between the level of RA-induced gene expression and the number of RA-induced ATAC-seq elements (Fig. 6E). Interestingly, 11% of RA-induced ATAC-seq peaks corresponded to RAR binding sites identified by ChIP-seq and possess the DR5 motif recognized by the RAR/RXR complex (Fig. 6D). This was the case for RA-induced peaks in the *dhrs3a, cyp26a1*/*b1* and *insm1b* genes (see red boxes in Fig. S6 and data not shown). However, many identified RAR sites did not display a significant increase in accessibility, such as those located in the *hoxba* locus (see green boxes in Fig. S6). Furthermore, the majority of RA-induced ATAC-seq peaks did not harbour RAR binding sites although they were usually found near RA-upregulated genes; instead, such peaks often harbour Gata or Hnf1b binding motifs (38% and 13% of all RA-induced elements, Fig 6D). Thus, it can reasonably be assumed that these genes are indirectly stimulated by RA signalling. For example, the pancreatic regulatory genes *pdx1, insm1a, rfx6* or *neurod1*, all upregulated by RA-treatment but devoid of RAR binding sites, had RA-induced ATAC-seq peaks located in their genomic neighbourhood and contained either Hnf1b or Gata binding motifs in such peaks (Fig. S7 and data not shown). To identify the GATA and Hnf1b family members involved in these indirect RA-regulations, we searched in the list of RA-stimulated genes (Table S2) for these 2 types of transcription factors and we also determined if RAR sites are detected in their genomic loci (Table S4 and S5). *hnf1ba* and *hnf1bb* were both induced by RA but the level of expression of *hnf1ba* was about 200-fold higher than *hnf1bb* in endodermal cells. Furthermore, *hnf1ba* contains three RAR binding sites located in evolutionary-conserved and nucleosome-free regions (Fig. 7A), while *hnf1bb* has only one weak RAR site (data not shown). Out of the 10 members of the GATA family, only *gata4* and *gata6* were significantly induced by RA but *gata6* was expressed in the endoderm at about 40-fold higher level compared to *gata4*. Furthermore, *gata6* has a high affinity RAR binding site in a nucleosome-free region located about 30 kb upstream from its TSS (Fig. 7B) while no such RAR site was present around the *gata4* gene (data not shown). Thus, all these analyses strongly suggest that, upon RA induction, RAR-RXR complexes directly activate expression of Gata6 and Hnf1ba, which in turn will open regulatory chromatin regions of many genes, including those coding for several pancreatic regulatory factors.

**Figure 7.**
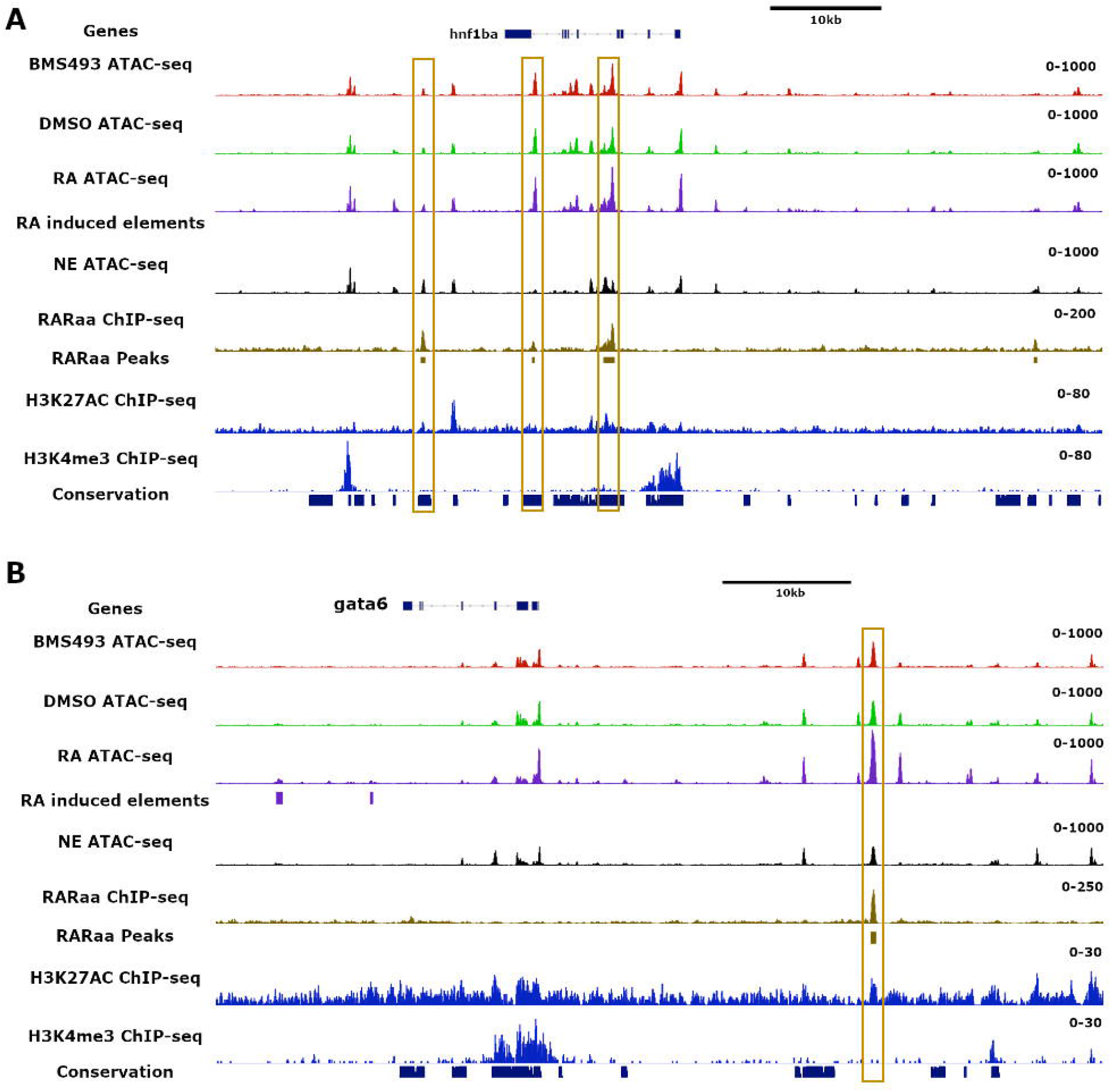
Location of RARa binding sites and of nucleosome-free regions in the *gata6* and *hnf1ba* gene loci. (A)Visualization of ATAC-seq reads from the merged 3 replicates obtained from endoderm treated with BMS493, DMSO or RA and from non-endodermal cells (NE), in addition to tracks showing RARaa binding sites, H3K27Ac and H3K3me3 marks determined by ChIP-seq. Regions showing conservation of genomic sequences among 5 fish species are also shown by blue boxes below the tracks. The RAR binding sites are highlighted by gold boxes.

## Discussion

In this study, we have used an integrative genomic approach to identify the genes regulated directly and indirectly by RA signalling in the endoderm at the end of gastrulation and notably involved in the specification of pancreatic progenitors. RNA-seq data performed on FACS-selected endodermal cells identified genes with enriched expression in endoderm and regulated by RA signalling. This gene set includes many genes previously shown to be regulated by RA in other germ layers like *hox, nr2f, cyp26a, dhrs3* or *hnf1b* genes (Feng et al. 2010b; C. E. Love and Prince 2012; Berenguer et al. 2020; Hernandez et al. 2004; Nolte, De Kumar, and Krumlauf 2019) indicating that the regulatory network triggered by RA is shared, at least partially, between the three germ layers. Analysis of the zebrafish RAR cistrome at the end of gastrulation confirms *hox* genes as direct RAR targets (Nolte, De Kumar, and Krumlauf 2019), acting on the anterio-posterior patterning of the endoderm, but the data also pinpoint 4 direct RAR target genes (i.e. *mnx1, hnf1ba, gata6* and *insm1b*) known to play a crucial role in pancreas development. Analysis of the nucleosome-free regions upon RA/BMS493 treatments reveals that, in addition to RAR, Gata and Hnf1b factors are the major transcription factors driving the increase in chromatin accessibility at numerous regulatory regions in endodermal cells. All together, these data indicate that Gata6 and Hnf1ba, which are directly regulated by RAR, activate in a second step the expression of many genes. A previous study on Xenopus explants showed the role of Hnf1b and Fzd4 downstream of RA signalling for the specification of pancreatic field (Gere-Becker et al. 2018). While our work confirm the conserved role of *hnf1ba* in zebrafish endoderm, we did not detect the involvement of zebrafish *fzd4* or other *fzd* paralogs which were not significantly induced by RA.

The treatment of embryos with exogenous RA leads to a similar number of up- and down-regulated genes; in contrast the treatment with the pan-RAR antagonist BMS493 mostly causes down-regulation of genes. Such difference can be explained by the mode of action of BMS394 which seems to stabilize the association of RAR with its co-repressors NcoR/Smrt (Germain et al. 2009). In contrast, RA causes mainly a release of co-repressors and the recruitment of co-activators. Most RA-downregulated genes detected in the present study likely represent indirect RAR targets such as genes expressed in anterior endodermal cells whose fate is modified upon RA treatment. This could be the case for the *nkx2*.*1, nkx2*.*3* and *tbx1* genes known to be expressed in the anterior endoderm and which are downregulated upon RA treatment. We also observed that the number of genes regulated by the antagonist BMS493 was much lower compared to the number of RA-regulated genes. Such a difference can be explained by the low proportion of endodermal cells receiving the endogenous RA signal. Indeed, BMS493 will affect only the cells under endogenous RA signalling (i.e. mid-trunk) while ectopic RA treatment affects all the other endodermal cells. In addition, a down-regulation is more difficult to detect for the genes having a low basal expression level or for genes expressed in a few endodermal cells.

In our study, we also determined the RAR cistrome at the end of zebrafish gastrulation to identify RAR direct targets. Comparison of the zebrafish RAR cistrome with the RAR cistrome of murine embryonic F9 cells (Chatagnon et al. 2015) identified 722 out of 2144 genes harbouring RAR sites in both zebrafish and mice, among which are several *Hox, Cyp26a, Meis* and *Nr2f* genes, as well as the 4 pancreatic regulatory genes *Mnx1, Gata6, Hnf1b* and *Insm1*. This conservation is further supported by the location of many RAR sites in CNEs revealing an evolutionary constraint to maintain these RAR sites. However, the majority of the zebrafish and murine RAR binding sequences could not be aligned. For example the two RAR sites located near the zebrafish *hnf1ba* and the murine *Hnf1b* genes are both located in fish and mammalian CNEs, respectively; these two RAR sites have also similar location in the murine and zebrafish genes (i.e. in the 4^th^ intron and downstream the Hnf1b gene) but the zebrafish and murine RAR sequences cannot be aligned. However, it is reasonable to assume that these RAR sites have a common origin and function.

Interestingly, 24 RAR sites were found in extremely conserved non-coding regions and sequences could be aligned from zebrafish to human. This is the case for RAR sites located in some *hox* clusters (Nolte, De Kumar, and Krumlauf 2019), near *meis2* (Berenguer et al. 2020), *nrip1a, ncoa3* and in the *skap1* gene (this study). Such high conservation suggests that the level of expression of these RA target genes must be precisely controlled. We show in the present study that the highly conserved RAR site in the 4^th^ intron of the zebrafish *skap1* gene drives expression in the neural tube and gut in a pattern reminiscent to *hoxb1b* expression, suggesting that this RARE controls the expression of the neighbouring *hoxbb* cluster. This is further supported by recent chromosome conformation capture (Hi-C) data obtained from adult zebrafish muscle tissue (Yang et al. 2020); mining these data show that the 4^th^ intron *skap1* CNE interacts with several *hoxbb* genes (not shown). Thus, this indicates that *hox* regulation by RA involved additional RAR sites located very far downstream from the *HoxB* cluster (i.e. more than 200 kb in the murine and human genome). This observation also reveals that the assignment of RAR sites to the closest neighbour could result in some incorrect links between RAR sites and their putative target genes. To improve RAR-gene assignment, ChIP-seq should ideally be combined with HiC on zebrafish gastrula. Still, we think that our strategy enabled to accurately define the majority of RAR-gene assignments as revealed by the correlation found between RA gene induction and the number of nearby RAR sites.

Our study also highlights that RA modifies the accessibility of thousands of genomic regions in zebrafish endodermal cells. The modest effect of BMS493 on chromatin is probably due to the small proportion of endodermal cells under the influence of endogenous RA, as already discussed above for the RNA-seq data. Interestingly, about 10 % of the sequences showing increased accessibility upon RA treatment correspond to RAR binding sites, while no RAR site shows reduced accessibility upon RA treatment. This observation supports the classical model where, upon RA binding, the RAR-RXR complex interacts with coactivators recruiting HAT which open chromatin structure (Niederreither and Dollé 2008). Still, many RAR sites do not display significantly increased chromatin accessibility despite strong activation of nearby genes by RA treatment. In such case, the RAR-RXR complex is often bound to a nucleosome-free region, even without RA signalling (i.e. upon DMSO- and BMS493-treatments), indicating that RAR-RXR regulates transcription through a mechanism other than changing chromatin structure. The increase of accessibility at 10% of RAR sites still suggests that the RAR-RXR complex contributes to the destabilization of nucleosomal structure at these sites and could act with (or as) pioneering transcription factors (Zaret 2020). Further experiments are needed to define more precisely the role of RAR-RXR in nucleosome removal.

About 90% of chromatin regions with increased accessibility upon RA treatment do not harbour RAR binding sites but rather contain GATA or HNF1b binding motifs. Both in zebrafish endoderm and in murine embryonic F9 cells, Gata6 and Hnf1b genes harbour RAR sites and are strongly induced by RA. Thus, these two factors seem to act as important effectors of RA signaling by opening chromatin at numerous regulatory regions. Gata6 and Hnf1b have previously been shown to be involved in pancreas development (Sun and Hopkins 2001; Haumaitre et al. 2005; Carrasco et al. 2012; Allen et al. 2011) and Gata factors have also been shown to act as pioneering factors (Zaret 2020; Bossard and Zaret 1998). Whether Hnf1b has an intrinsic ability to bind nucleosomal DNA and to open chromatin or whether this requires the involvement of other co-factors remain to be determined.

By determining the transcriptomic landscape and genome-wide chromatin accessibility of endodermal and non-endodermal cells at the end of gastrulation, the present data also provide the lists of genes and regulatory cis-elements specifically active in the endoderm. For example, the RAR site located downstream from the *mnx1* gene is located in a region which is nucleosome-free only in the endoderm. Accordingly, our transgenic reporter assay shows that this region drives expression in the posterior endoderm and in the dorsal pancreas. Other endoderm-specific ATAC-seq peaks showed similar endoderm-specific activity (data not shown). Thus, our ATAC-seq data and RNA-seq data allows the genome-wide identification of endoderm specific enhancers. This strategy can also help to assign a regulatory region to its target gene by correlating expression and chromatin structure; the accuracy of such analyses can be increased by including more data from different cell types or from different developmental stages.

In conclusion, the present study provides interesting resources not only to decipher the regulatory network triggered by RA but also to unveil regulatory regions involved in endoderm formation and patterning in zebrafish.

## Material and methods

### Zebrafish strains, sample preparation and cell purification by FACS

Fish were maintained in accordance with the national guidelines and all animal experiments described herein were approved by the ethical committee of the University of Liege (protocol number 1980). Endodermal and non-endodermal cells were obtained using the transgenic line Tg(*sox17*:GFP) (Mizoguchi et al. 2008). Transgenic embryos were incubated with DMSO (0.1% volume) as control, 1 μM RA or 1 μM BMS943 from 1.25 to 11 hpf. Embryos were analysed either at 11hpf (3-somite stage) or either washed in embryos medium, raised until 13.5 hpf (8-somite stage) in embryos medium. Embryos were then dechorionated, deyolked, dissociated in FACS solution (HBSS without Ca++ and Mg++ and containing 1% BSA) by gentle pipetting and cells were directly sorted by FACS Aria II based on the GFP expression. Sorting was performed as a single run in purity mode and three replicates were prepared for each treatments. The efficiency of RA and BMS493 treatments were verified on Tg(pax6:GFP) embryos through the direct observation of fluorescent ectopic pancreatic endocrine cells in anterior endoderm (for RA treatment) and the absence of endocrine pancreatic cells (for BMS493 treatment).

### RNA-seq library preparation and data analysis

cDNA was obtained using the SMART-seq2 protocol as recently described (Lavergne et al. 2020). Briefly, each cell preparation was pelleted and lysed in Lysis Buffer by freezing in liquid nitrogen and stored at -80°C. Synthesis of cDNA was performed directly on the lysed cells; cDNA were amplified by 10 PCR cycles and purified before assessing their quality on the bioanalyzer (2100 high sensitivity DNA assay, Agilent Technologies). 150 pg of cDNA were used as input to prepare the libraries using the Nextera XT DNA kit (Illumina). 75 bp single-end sequences were obtained using the NextSeq500 Illumina Sequencer with coverage of about 20 million reads per library.

Raw reads were aligned to the zebrafish genome (Zv9, Ensembl genome version 79, ensembl.org) using STAR software (Dobin et al. 2013). Normalization and differential expression analysis were performed using DESeq2 (M. I. Love, Huber, and Anders 2014). Genes were considered differentially expressed with FDR < 0.01 (False Discovery Rate).

### Expression of RARaa-myc in zebrafish embryos

The whole zebrafish RARaa coding sequence was amplified by RT-PCR and inserted into the pCS2MT vector in frame with Myc-epitope tags located at the C-terminal end. The Myc-RARaa mRNA was synthesized from this plasmid by *in vitro* transcription (mMESSAGE mMACHINE sp6 transcription kit, Invitrogen) and injected into fertilized zebrafish eggs. Validation of Myc-tagged RARaa protein expression was done by western-blotting on cytosolic and nuclear lysate fractions from injected embryos using a ChIP-grade anti-myc antibody (ab9132, Abcam).

### ChIP-seq library preparation and data analysis

About 0.5 nL of myc-tagged RARaa mRNA was injected in zebrafish fertilized eggs at a concentration of 70 ng/µL and embryos were incubated until they reached bud stage (10.5 hpf, end of gastrulation). RA was added to the medium at a final concentration of 1 μM and embryos were grown for an extra hour. Around 2000 injected embryos were used for chromatin immunoprecipation essentially as previously described (Morley et al. 2009) and fixed with 1.85% PFA for 10 minutes. Diagenode Bioruptor sonicator was used for chromatin shearing. Dynal Protein A magnetic beads (Diagenode) and ChIP-graded anti-myc antibody (ab9132, Abcam) were used for the immunoprecipitation, an aliquot of chromatin being taken before the immunoprecipitation step and used as negative control. Libraries were prepared with the NEBNext Ultra II DNA Library Prep kit (Bioke). 42 bp pair-end sequences were obtained using the NextSeq500 Illumina Sequencer with coverage of 60 million reads per library.

Raw reads were mapped to the zebrafish genome (Zv9) using bowtie2 (Langmead and Salzberg 2012) using the settings “--threads 3 --very-sensitive”. Enriched peaks were called using MACS2 (Zhang et al. 2008) with the following settings: callpeak --gsize 1.4e9-- nomodel --call-summits --qvalue 0.05. Motif enrichment analysis was performed using HOMER (Heinz et al. 2010) adding a motif length from 6 to 18 bp, to ensure the inclusion of the different RARE. Annotation of peaks was done using ChIPseeker (Yu, Wang, and He 2015).

### ATAC-seq library preparation and data analysis

After isolation by FACS, cells were pelleted and ATAC-Seq libraries were prepared as described elsewhere (Buenrostro et al. 2015). Libraries were sequenced 42 bp paired end using the NextSeq500 Illumina Sequencer with coverage of 40 million reads per library. Three replicates were prepared for each condition.

Raw reads were mapped to the zebrafish genome (Zv9) using bowtie2 (Langmead and Salzberg 2012) with the following settings: --threads 3 --very-sensitive --maxins 2000 --no-discordant. ATAC-seq peaks were identified using MACS2 (Zhang et al. 2008) with the settings: --nomodel --shift 40 --extsize 80 -q 0.05 --gsize 1.4e9. Histone modification peaks were obtained from available datasets (GEO accession number GSE48254)(Paik et al. 2013) as well as the zCNEs (Hiller et al. 2013). Quality assessment of ATAC-seq libraries was performed with ATACseqQC (Ou et al. 2018). ChIPpeakAnno (Zhu et al. 2010) was used to create the density and heatmap plots. Enriched peaks for ATAC-seq datasets were called using MACS2. Peaks showing different signal among ATAC-seq samples were identified with DiffBind (Ross-Innes et al. 2012). Briefly, the density of mapped reads was calculated for each of the 156604 regions obtained by merging the peaks obtained for each of the 12 individual samples and differential analysis was performed in these regions to identify the regions with differentialy opened chromatin in each condition. Motif enrichment analysis for each set of treatment specific peaks was performed using HOMER (Heinz et al. 2010). Annotation of peaks was done using the ChIPseeker (Yu, Wang, and He 2015).

### Data integration and visualization

To study the correlation of ATAC-seq/ChIP-seq and RNA-seq a student’s t-test was performed to compare the read counts in RNA-seq of each gene and the transcription start site (TSS) accessibility in ATAC-seq/ChIP-seq. Correlation of flanking treatment/cell specific and common ATAC-seq elements, as well as, ChIP-seq peaks and log2 fold change from RNA-seq was analysed by Spearman’s rank correlation.

### Generation of zebrafish transgenic reporter lines

The sequences containing the RAR binding sites downstream from the mnx1 gene and in the 4^th^ intron of skap1 gene were amplified by PCR (primers for mnx1: AATGTACAATTGATCCCATTCGG and TAAACTTTATCACTGTGTCAGATCA; primers for skap1: AGGGGAAATTGACTGTGTCTTGCT and TCAACCAGGCCATTCCAAGTGA). These fragments of respectively, 441bp and 812 bp, were inserted in topo-GW plasmid and transferred upstream of the c-fos minimal promoter followed by EGFP coding sequences (pGW-cfosEGFP plasmid)(Fisher et al. 2006) by gateway LR recombination. The purified RAR-mnx1:GFP and RAR-skap1:GFP transgene flanked by Tol2 inverted repeats were injected into 1–2-cell zebrafish embryos with Tol2 transposase mRNA. Embryos presenting GFP expression were raised to adulthood and crossed to generate stable transgenic lines. For both transgenes, several lines were analyzed and showed similar expression pattern indicating that expression results from the inserted RAR fragment and not from the transgene insertion sites. GFP expression from the stable transgenic embryos were analysed with a Leica fluorescence stereomicroscope.

## Declarations

All animal experiments were conducted according to national guidelines and were approved by the ethical committee of the University of Liège (protocol numbers 1980). The raw datasets generated in the current study will be deposited on ENA (in progress). The authors declare that they have no competing interests.

## Acknowledgments

We thank the following technical platforms of GIGA from Liège university: GIGA-Zebrafish (H Pendeville), GIGA-Cell Imaging and Flow Cytometry platform (S Ormenese and S Raafat), GIGA-Genotranscriptomic (W Coppieters, and L Karim). We are indebted to Virginie Vonberg for technical help in transgene constructions and for zebrafish egg injections. We also thank Fabio D’Orazio and Nan Li for assistance with the ATAC-seq protocol. We acknowledge all members of the ZENCODE-ITN project for helpful discussions and feedback.

## Competing interests

No competing interests declared.

## Funding

This work was supported by ZENCODE-ITN European project 643062 (A.R.L., B.L., F.M., F.C.W. and B.P.) and by the Léon Fredericq fund (A.R.L. and B.P.). B.P., I.M., and M.L.V. are associate researchers from FRS/FNRS (Fonds National pour la Recherche Scientifique).

## Authors’ contributions

A.L.P. carried out the RNA-seq, ChIP-seq and ATAC-seq experiments and performed the bioinformatic analyses with help from P.B. and B.L.. F.M. and F.W. helped in the setting of ATAC and ChIP experiments and for funding acquisition. B.P., A.L.P., M.L.V. and I.M. conceived the study, interpreted the data and wrote the manuscript. All authors read, edited and approved the final manuscript.

## Data availability

All data generated and analysed in this study are available in the supplemental tables. The raw RNA-seq, ChIP-seq and ATAC-seq data will be deposited on Gene Expression Omnibus (GEO).

## Notes

### Competing Interest Statement

The authors have declared no competing interest.

