## Supplementary material for "Identification of downstream effectors of retinoic acid specifying the zebrafish pancreas by integrative genomics": SuplementalData

**<u>Supplementary Figures :</u>**

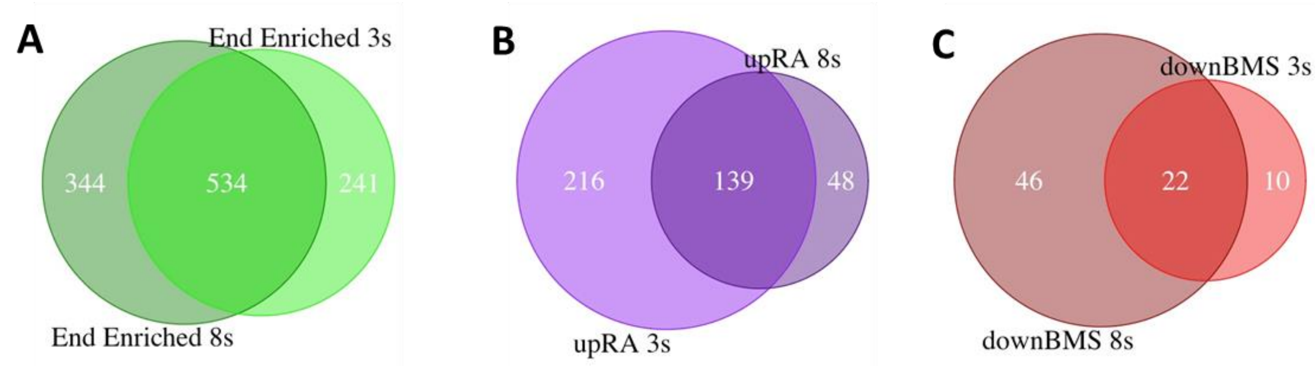


**Suppl. Figure 1 : Overlap in the sets of genes showing endoderm-enriched expression and regulation by RA signalling at 3- and 8- somites stages.**
Venn diagrams revealing the overlap in the set of genes with endodermal enriched expression (A), up-regulated by RA (B) and down-regulated by BMS493 (C) at 3- and 8-somites stages.

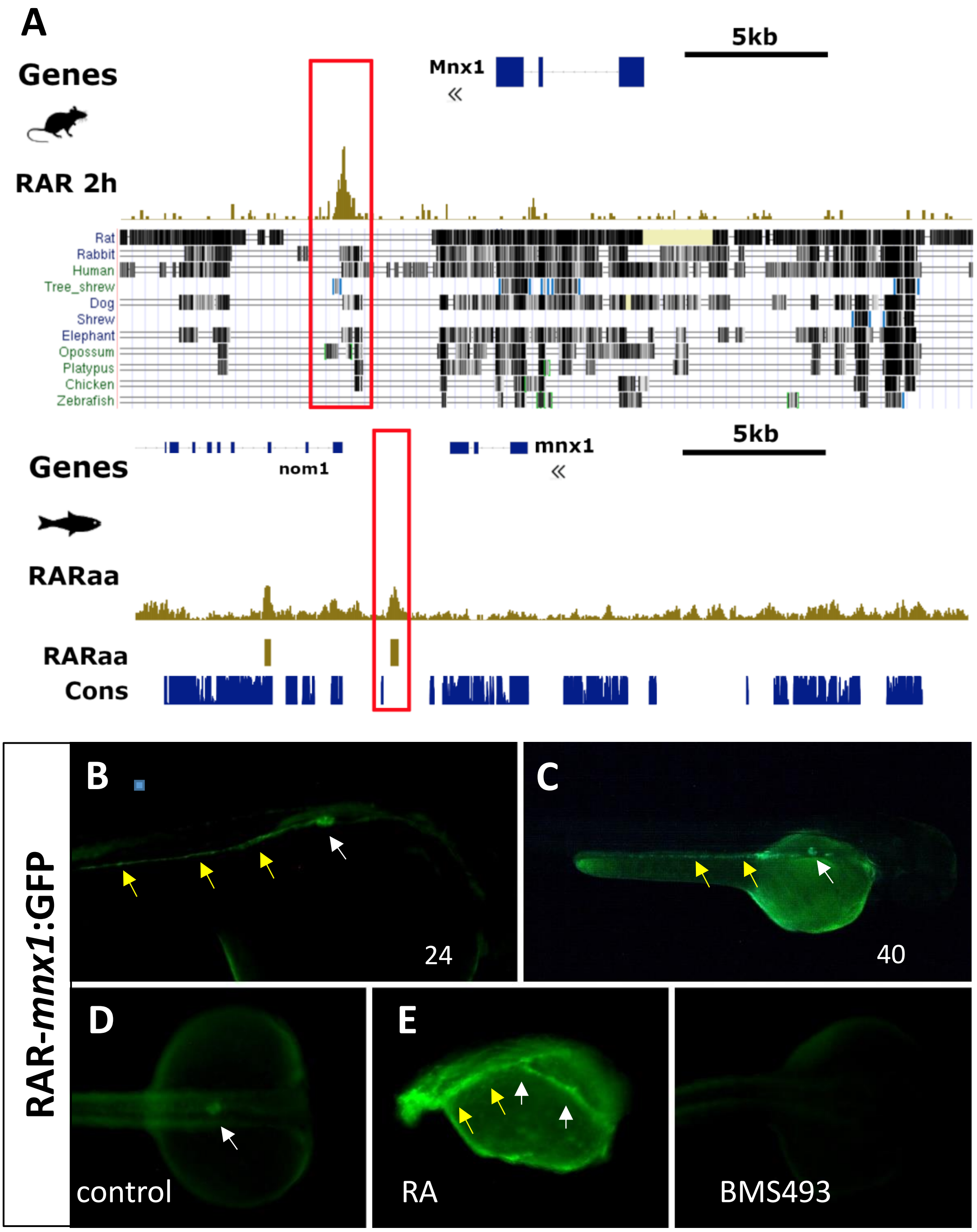


**Suppl. Figure S2. A RAR binding site is located at an equivalent place downstream the zebrafish and murine mnx1 gene.**
(A) Genome browser views of the Mnx1 gene locus in mouse (upper panel) and in zebrafish (lower panel) showing the presence of a RAR binding site downstream the gene by ChIP-seq analysis (gold tracks) and located at the border of a conserved regions. Sequence conservation is highlighted below the murine ChIP-seq peaks through the comparison with several vertebrate species (data from UCSC genome browser) as well as by the fish conserved genomic regions ("Cons. track" in blue). (B-F) The mnx1 RAR element is able to target GFP reporter expression in the dorsal pancreas (white arrow) and in the posterior gut (yellow arrows) in zebrafish transgenic assays at 24hpf (B) and 40hpf (C) stages. GFP expression is slightly increased by RA (panel E) and switch off by BMS493 (panel F) (treatments being performed during gastrulation).

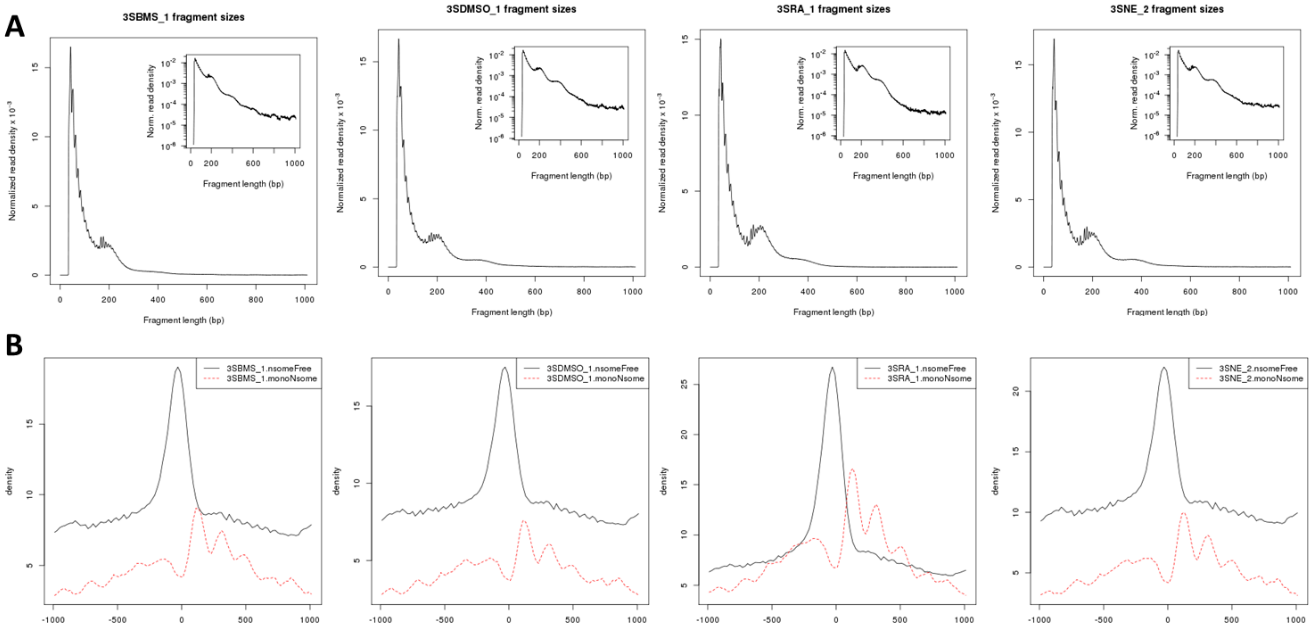


**Suppl. Figure S3. Distribution of ATAC-seq fragments**

(A) Fragment length distribution obtained by ATAC-seq for one replicate of each condition : BMS493-, DMSO-, RA-treated endodermal cells and non-endodermal cells. Fragments with size below of 180 bp derive from nucleosome-free regions; the second peak with length size of about 200 bp derive from mononucleosome region. (Inset) log-transformed histogram showing monomer and dimer of nucleosomes. (B) Distribution of reads around ENSEMBL-annotated transcription start sites (TSS, centered at 0) for nucleosome-free fragments (i.e. <180bp)(black line) and for mono-nucleosome fragments (i.e. +-200 bp)(red line). As expected, sequences just upstream TSS are enriched in nucleosome-free fragments while mononucleosomes lie downstream TSS with a periodicity.

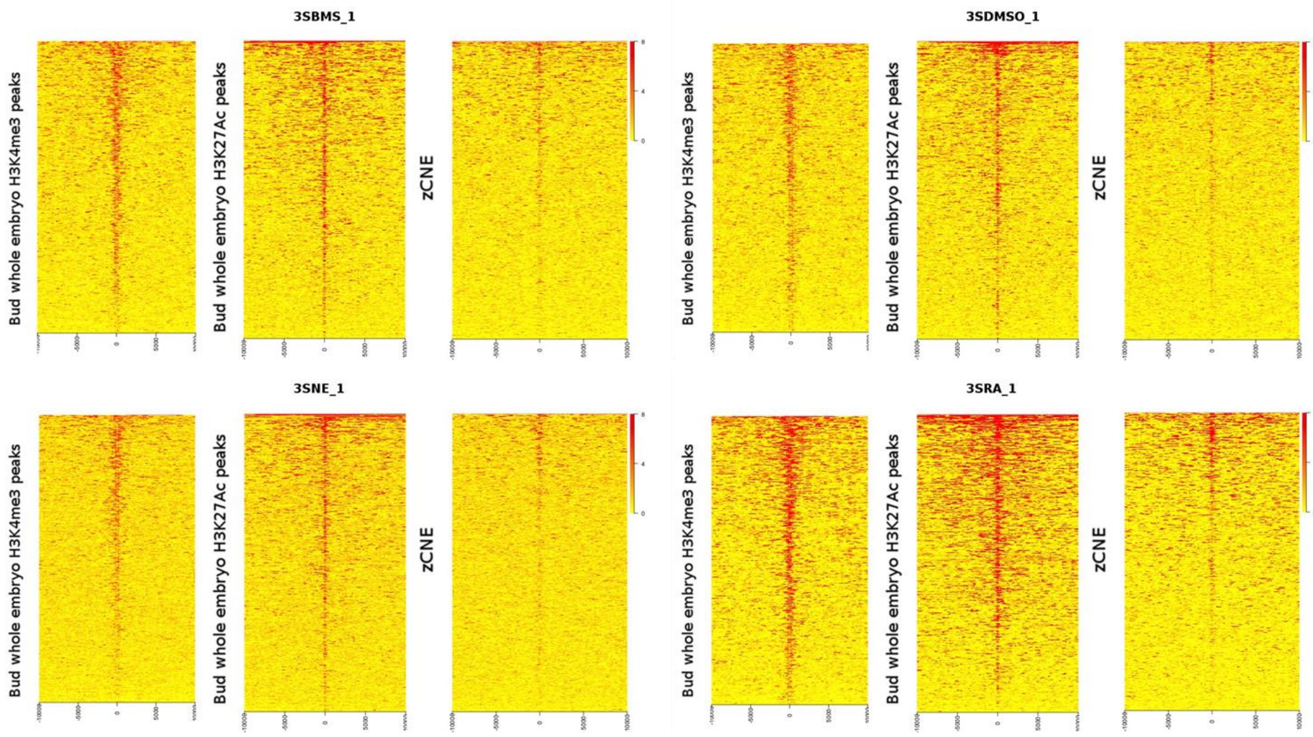


**Suppl. Figure S4. ATAC-seq framents display hallmarks of regulatory regions.**

Heat-maps showing enrichment of ATAC-seq reads at the middle of chromatin regions harbouring H3K4me3 and H3K27Ac epigenetic marks and at genomic areas corresponding to fish conserved non-coding elements (zCNE). The maps display intervals flanking 10 kb up and downstream of the features. The heat-map plots display the analysis for the 12 ATAC-seq samples obtained at 3-somites stage with endodermal cells treated with BMS493 (3SBMS), with DMSO (control)(3SDMSO), with RA (3SRA) and obtained with non-endodermal cells (3SNE).

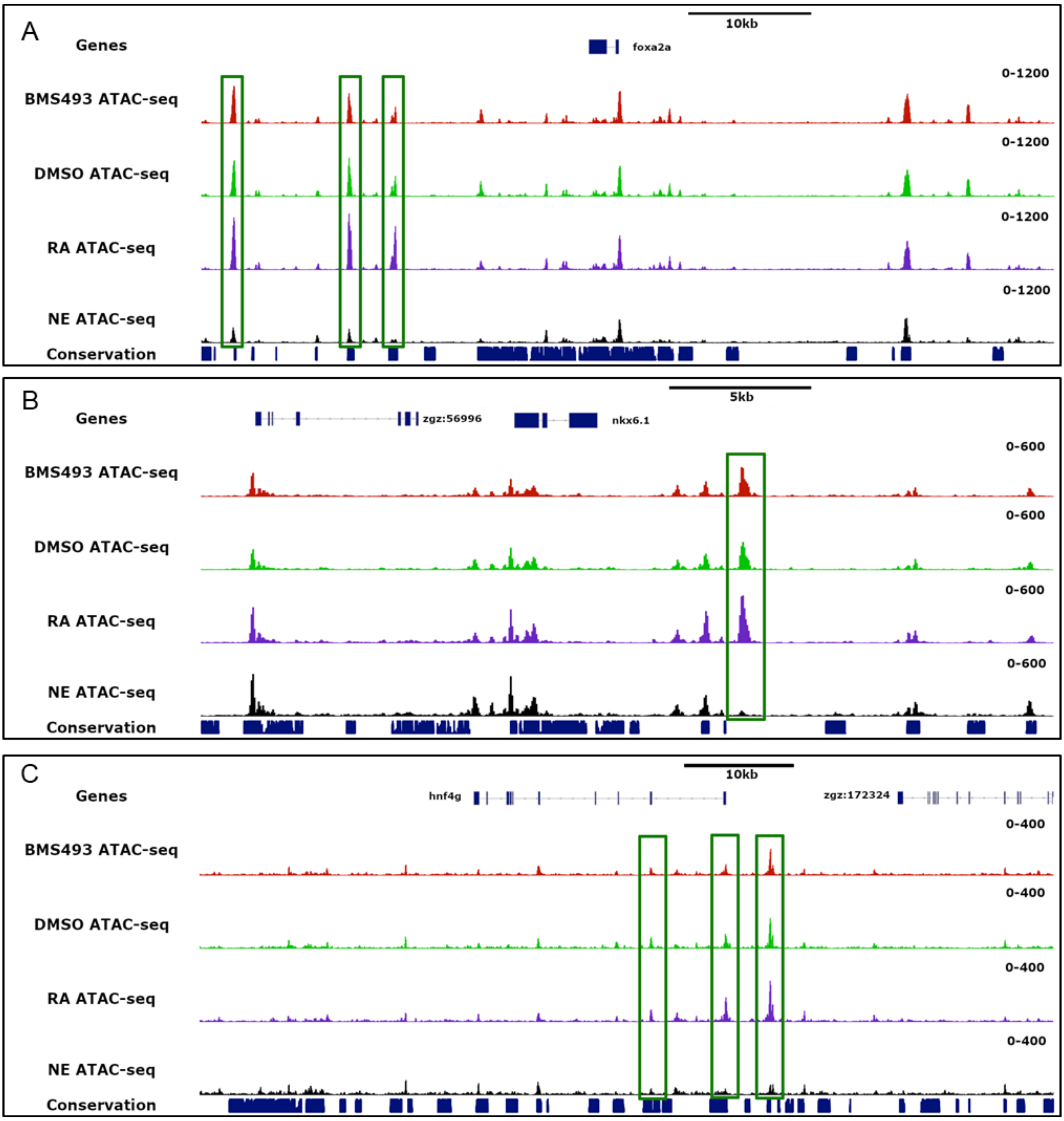


**Suppl. Figure S5: Presence of endoderm-specific ATAC-seq peaks in genomic loci harbouring endoderm and pancreatic regulatory genes.**

Gene browser views of the *foxa2* (A), *nkx6.1* (B) and *hnf4g* (C) genomic loci displaying ATAC-seq peaks obtained from endoderm (track1 : treated with BMS493), (track2 : treated with DMSO ), (track3: treated with RA ) and non-endodermal cells (track4 : NE treated with DMSO). Tack5 displays conserved genomic sequences (phastcons). The ATAC-seq peaks shown in green boxes are endoderm-specific.

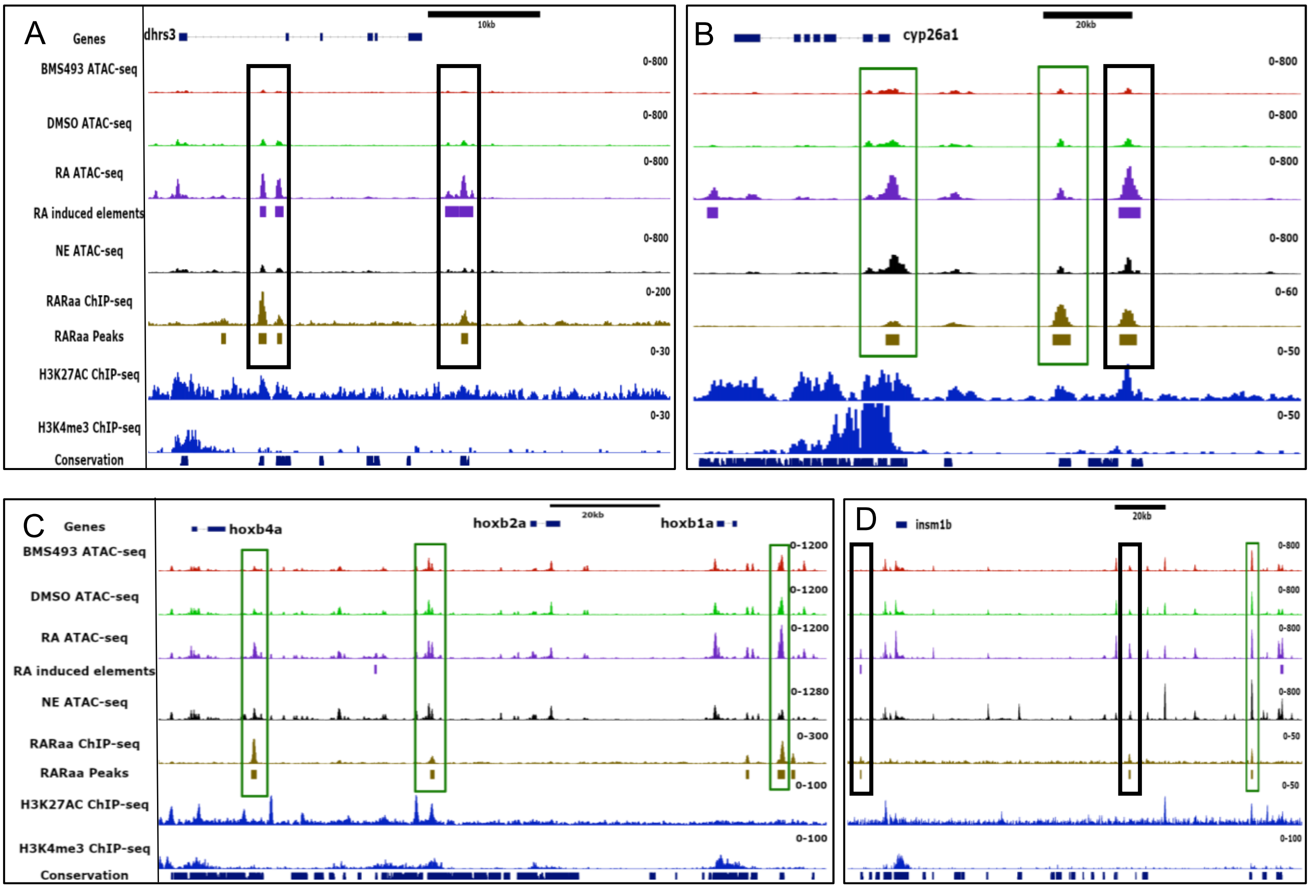


**Suppl. Figure S6. Chromatin accessibility is increased at some RAR binding sites upon RA treatment.**

Gene browser views of the direct RAR target genes *dhrs3a* (A)*, cyp26a1* (B), *hoxb1a* (C) and *insm1b* (D). Each panel displays the ATAC-seq peaks from endoderm treated with BMS493 (track1), with DMSO (track2) and with RA (track3) and non-endodermal cells (NE)(track 5). RARaa ChIP-seq (track 6), H3K27Ac (track8) and H3K4me3 (track9) ChIP-seq peaks are indicated. The RA induced elements from ATAC-seq and RARaa ChIP-seq are shown in tracks 4 and 7, respectively. The RAR binding sites which are more accessible upon RA treatment are shown by red boxes (FDR<0,05). The green boxes highlight the RAR sites whose chromatin accessibility is not significantly modified by RA treatment (FDR>0,05).

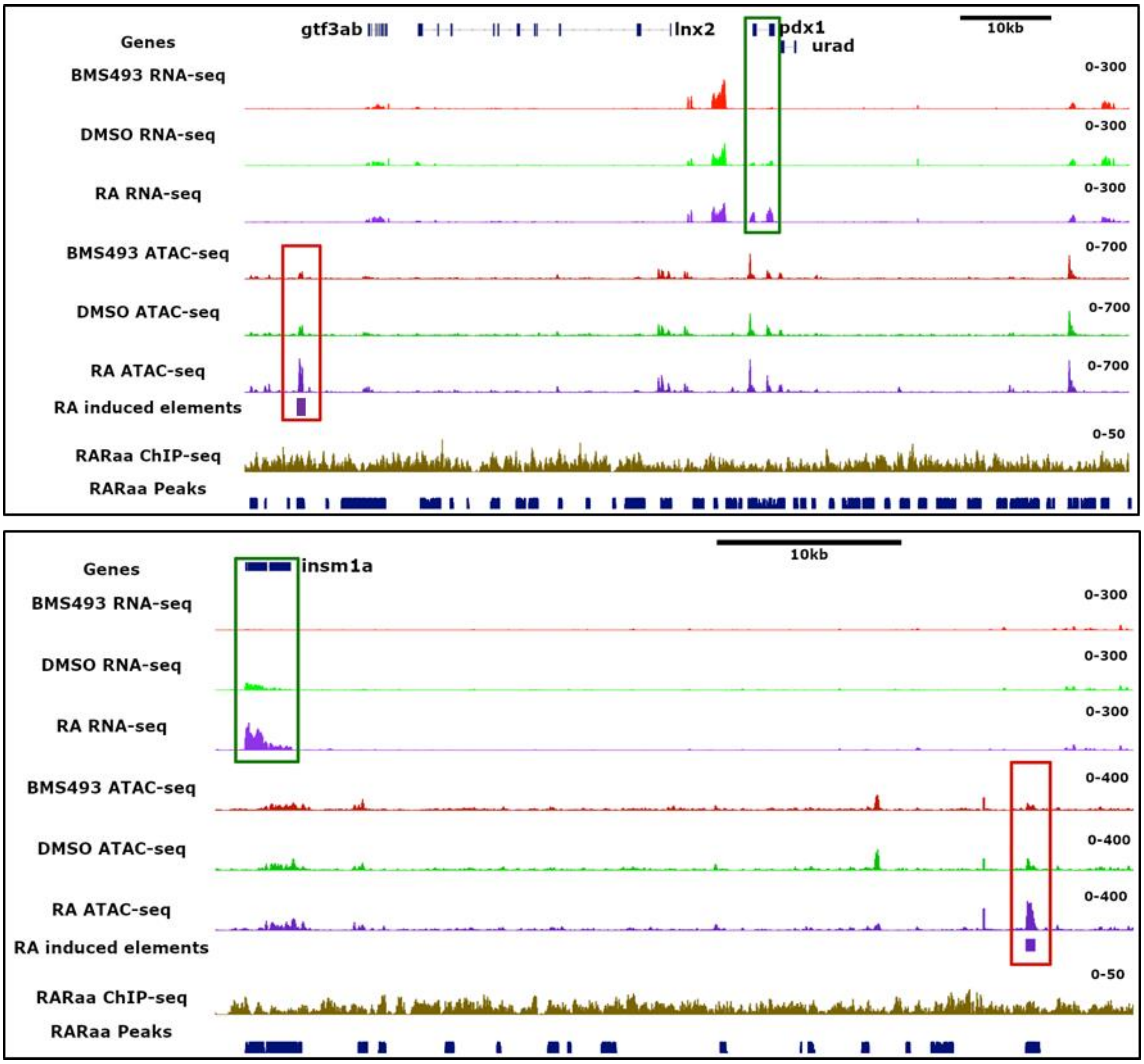


**Suppl. Figure S7 : Nucleosome-free regions induced by RA treatment near the pdx1 and insm1a genomic loci.**

The first 3 tracks show RNA-seq reads from endoderm treated by BMS493 (track1), by DMSO (track2) and by RA (track3). The green boxes highlight the increase of pdx1 mRNA (upper panel) and insm1a mRNA (lower panel). The tracks 4 to 7 show the ATAC-seq peaks from endoderm treated by BMS493 (Track4), by DMSO (track5) and by RA (track6). Track 7 indicates the peaks induced by RA treatment and track8 shows absence of RARaa ChIP-seq peak. The red boxes highlight the peaks induced by RA treatments. The two RA-induced elements display sequence conservation as shown in the phastCons track (track9)(blue boxes).

**Supplementary Tables :**

**Suppl. Table S1: List of genes having significant enriched expression in endodermal cells at 3-somites stage (excel sheet 1) and at 8-somites stage (excel sheet 2).**

The enrichment is given by log2 of fold change (FC)(RA versus DMSO) with the fold discovery rate (FDR). Normalized level of expression is given in count of reads per million (CPM) for the DMSO and RA treatments. Enriched genes were selected with a FDR<0.01.

**Suppl. Table S2: List of genes regulated by RA treatments at 3-somites stage (excel sheet 1) and at 8-somites stage (excel sheet 2).**

The enrichment is given by log2 of fold change (FC)(RA versus DMSO) with the fold discovery rate (FDR). Normalized level of expression is given in count of reads per million (CPM) for the DMSO and RA treatments. RA-regulated genes were selected based on FDR <0.01.

**Suppl. Table S3: List of genes regulated by BMS493 treatments at 3-somites stage (excel sheet 1) and at 8-somites stage (excel sheet 2).**

The enrichment is given by log2 of fold change (FC)(BMS493 versus DMSO) with the fold discovery rate (FDR). Normalized level of expression is given in count of reads per million (CPM) for the DMSO and BMS493 treatments. BMS493-regulated genes were selected based on FDR <0.01.

**Suppl. Table S4: List of RAR binding sites identified by ChIP-seq.**

Location of all 4750 RARa ChIP-seq peaks is  given by their coordinates (start, end and width of peaks) on each zebrafish chromosome.

**Suppl. Table S5: List of genes regulated by RA treatment and harbouring a RARa sites in their genomic vicinity.**

Location of ChIP-seq peaks and the associated genes is given by their coordinates on each chromosome. Gene identity is given by their Ensembl ID, transcript ID and gene name. The location of the RAR ChIP-seq peaks and the distance from their assigned gene is given in columns I and J, respectively. Sites assigned to a gene stimulated by RA are given in excel sheet1; RAR sites assigned to a repressed gene are given in excel sheet 2.

**Suppl. Table S6: List of RARa binding sites located in zCNE.**

Location of zebrafish (excel sheet 1) and murine (excel sheet 2) RARa sites which are located in evolutionary conserved genomic sequences, with their assigned gene (given by Ensembl ID and gene name). Coordinates of ChIP-seq peaks (columns A-D), their location and distance to the assigned gene TSS are given in column I and J. The RARa sites located in highly conserved sequences from fish to mice are indicated in column L (excel sheet 1).

**Suppl. Table S7: List of ATAC-seq peaks specifically identified in endodermal or non-endodermal cells.**

Location of ATAC-seq peaks identified specifically in endoderm (sheet 1) or non-endoderm (sheet2) is given by their coordinates (start, end and width of peaks) on each chromosome. Peaks enriched in each cell type were selected based on FDR<0.05.

**Suppl. Table S8: List of ATAC-seq peaks affected by RA treatments.**

Location of ATAC-seq peaks induced (sheet 1) and repressed (sheet 2) by RA treatments is given by their coordinates on each chromosome. The gene assigned for each peak is given by their Ensembl ID and name. Selection of peaks was done based on FDR<0.05 (RA versus BMS493).
